# Proteomics-Based Discovery of Symmetry-Specific Readers and Antireaders of 5-Formylcytosine in Mammalian DNA

**DOI:** 10.64898/2026.04.09.717454

**Authors:** Zeyneb Vildan Cakil, Lena Engelhard, Nina Seidel, Simone Eppmann, Tanja Bange, Daniel Summerer

## Abstract

5-methylcytosine (mC) and its oxidized derivatives are epigenetic modifications of mammalian DNA that play key roles in transcription, cell differentiation, and cancer. They predominantly occur within palindromic CpG dyads, creating multiple possible combinations across the two dyad strands, each representing a chemically distinct DNA major groove mark. Among the modifications, 5-formylcytosine (fC) interacts with a large number of proteins and has been associated with important roles in chromatin regulation. However, it is poorly understood whether nuclear proteins selectively recognize the symmetry of individual fC-containing CpG dyads. Here, we report the first proteome-wide interaction profiles of symmetric and asymmetric fC modifications in mammalian DNA. Our analysis spans several human and mouse cell lines and three DNA promoter probes containing CpG dyads with five distinct combinations of fC, C, and mC nucleobases. We identify a diverse set of fC reader proteins with distinct symmetry preferences, including transcription factors (e.g., MAX, HEY1, RFX5, SIX1, SIX2, and FOXJ3) and DNA repair proteins (MPG and TDG). Notably, some proteins act as fC readers or anti-readers depending on the sequence context, suggesting a role for fC in modulating their target selection. Our data reveal widespread symmetry-specific recognition of fC by mammalian proteins and provide a rich resource for studying its potential functions in chromatin regulation.

**GRAPHICAL ABSTRACT:** 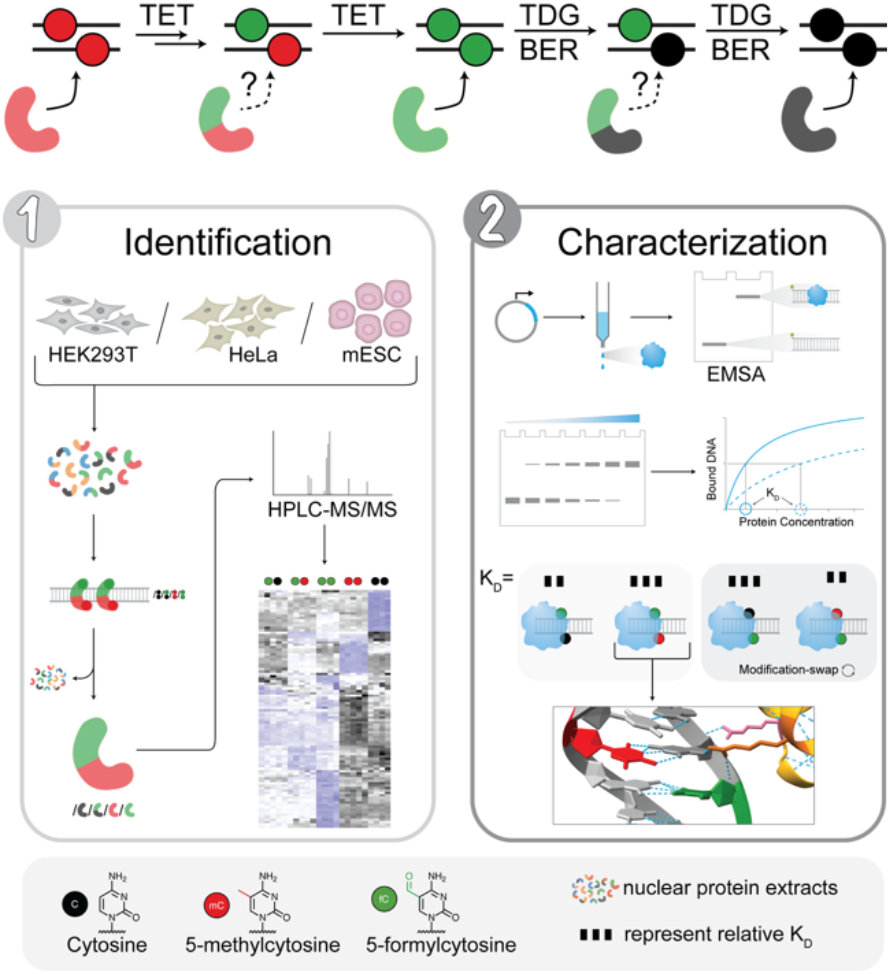

## INTRODUCTION

5-methylcytosine (mC) is the central epigenetic mark of mammalian DNA and regulates differentiation, development, and carcinogenesis (1,2). 5mC is introduced and maintained by DNA methyltransferases (DNMTs) primarily in palindromic CpG dyads, 60–80% of which are methylated in somatic cells (3,4). mC can be removed passively over cell-divisions, but also be oxidized to 5-hydroxymethylcytosine (hmC), 5-formylcytosine (fC), and 5-carboxylcytosine (caC) by ten-eleven-translocation (TET) dioxygenases (5,6), providing the possibility of active demethylation via thymine-DNA glycosylase (TDG) excision and base excision repair (BER) (**Figure 1a**) (7,8). Among the three oxidized mC derivatives, fC has been attributed unique properties and functions. For example, it increases DNA flexibility, which plays a role in nucleosome stabilization (9,10). The electrophilic 5-formyl group equips fC with the unique ability to form transient imine bonds with protein lysines (11,12), which has been implicated to play a role in nucleosome positioning (13). Moreover, fC impairs RNA polymerase II elongation and fidelity (14), and during *Xenopus* zygotic genome activation, it acts as an instructive mark by promoting RNA polymerase III recruitment and tRNA expression (15). Unravelling additional functions of fC is closely connected to a system-wide understanding of how it interacts with nuclear proteins. Indeed, previous mass spectrometry (MS)-based proteomics studies employing oxi-mC-modified DNA enrichment probes have generally identified a larger number of readers for fC than for hmC and caC and provided new clues to possible functions (16-18). However, a central aspect of fC-protein interactions remains poorly understood: The co-existence of mC and “oxi-mC” in palindromic CpG dyads can theoretically lead to fifteen different symmetric and asymmetric dyad states (that is, combinations of cytosine forms across the two CpG strands (19) (**Figure 1b**), each representing a unique physicochemical mark in the DNA major groove with potentially unique roles in chromatin regulation (20). Previous proteomics studies have focused exclusively on symmetric fC CpG dyads (“fC/fC”) (16-18). Proteomics studies with other prevalent fC dyads–conducted in a comparative fashion– are therefore urgently needed to obtain a refined picture of fC interactomes, that is, to reveal actual fC dyad-specificities of proteins, and to provide clues to potential symmetry-dependent regulatory functions. Indeed, symmetry-dependent interactomes have been observed in comparative proteomics studies focusing on hmC (21), and differences between symmetrically and hemi-modified fC-CpG dyads have been described for recombinant transcription factors (22).

**Figure 1.**
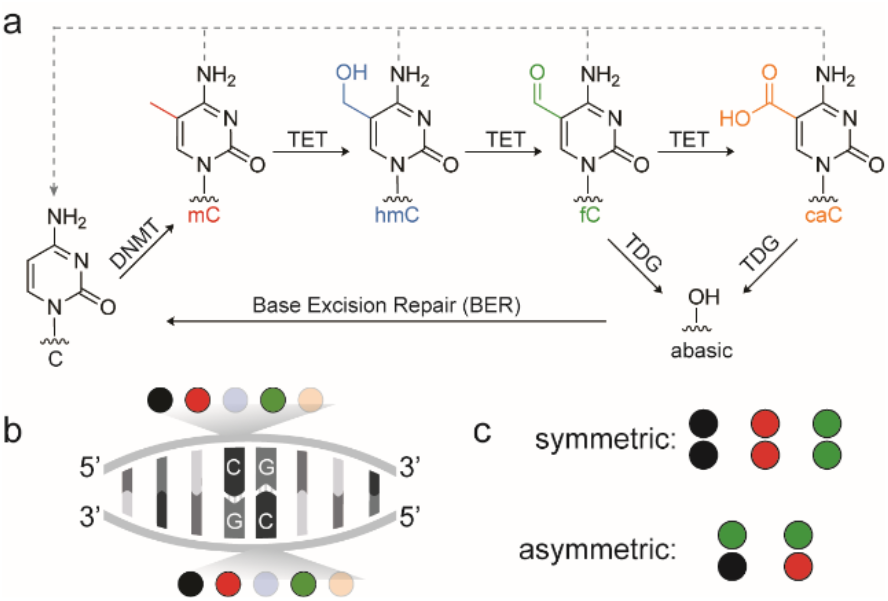
Epigenetic cytosine modifications in CpG dyads. (a) Cytosine methylation and passive (dashed lines) or active (solid lines) demethylation in mammals. DNMT: DNA methyltransferase, TET: ten-eleven translocation dioxygenase, TDG: thymine DNA glycosylase. (b) Combinations of cytosine 5-modifications in CpG dyads. (c) Strand-symmetric and -asymmetric combinations of C, mC and fC in CpG dyads evaluated in this study. The modification colour code is used throughout the manuscript.

We here report comparative proteome-profiling studies for five symmetric and asymmetric CpG dyads, i.e. C/C, mC/mC as well as fC paired with C, mC and itself (**Figure 1c**). Our study covers two different human cell lines, mouse embryonic stem cells (mESC), as well as three different DNA probe designs derived from human and mouse promoters. We observe distinct binding preferences of many proteins towards individual fC-CpG dyad states. Reader and antireader proteins interact with fC-CpG dyads in a sequence (probe)-specific manner and are highly organism- and tissue-specific. For all probes and tissues, we find a similar number of readers for fC dyads as for the canonical C/C and mC/mC dyads combined. Moreover, the three fC dyads states generally attract very different reader groups and also significantly differ from the reader groups of C/C and mC/mC dyads. Further characterization of selected readers confirms distinct preferences for specific fC CpG dyad states for transcription factors (TFs; MAX, HEY1, RFX5, SIX1, SIX2, FOXJ3) and BER proteins (MPG and TDG). We also observe for some proteins that they read or antiread specific fC dyads depending on the sequence context, hinting at a complex interplay between modifications and protein’
ss sequence selectivities. Our data provide biochemical support for the view that functions of fC are symmetry-dependent, and they provide a rich resource for studies aiming at unravelling regulatory mechanisms in chromatin controlled by symmetric and asymmetric fC modifications.

## MATERIAL AND METHODS

### Cell Culture

HEK293T and HeLa cells were cultivated in DMEM (Dulbecco’s Modified Eagle Medium) (PAN) supplemented with 10% FBS (PAN), 2 mM L-glutamine (PAN), 100 U/mL penicillin and 0.1 mg/mL streptomycin (PAN) in humidified incubator (≥ 95%) at 37°C and 5% CO_2_. mESC (E14TG2a) were cultured on 0.1% gelatin-coated tissue culture flasks in serum+LIF medium composed of GMEM (Glasgow’s Minimum Essential Medium) (Thermo Fisher Scientific), 10% ES-qualified, batch tested FBS (Thermo Fisher Scientific), 1x non-essential amino acids (Thermo Fisher Scientific), 2 mM GlutaMAX (Thermo Fisher Scientific), 1 mM sodium pyruvate (Thermo Fisher Scientific), 0.1 mM 2-mercaptoethanol (Thermo Fisher Scientific), 100 ng/ml LIF (Max Planck Institute protein expression facility) and 100 U/mL penicillin and 0.1 mg/mL streptomycin (Thermo Fisher Scientific) in humified incubator (≥ 95%) at 37°C and 5% CO_2_.

### Nuclear Extraction

The nuclear proteins were extracted from HEK293T, HeLa and mESC cultures as described previously(21). Briefly, cells were washed with DPBS and detached using trypsin for 3-5 minutes at 37 °C. Cells were harvested by centrifugation at 300 g for 5 minutes. The resulting pellet was washed twice with ice-cold DPBS and then resuspended in ice-cold cell lysis buffer (10 mM HEPES, pH 7.9; 10 mM KCl; 1.5 mM MgCl_2_; 0.1 mM EDTA; 0.5 mM DTT; 0.5 mM PMSF) supplemented with a protease inhibitor cocktail (Roche). After a 10-minute incubation on ice, 0.5% IGEPAL was added, and the mixture was vortexed. The sample was then centrifuged at 3000 g for 20 minutes at 4 °C. The pellet was washed twice with cell lysis buffer. The pellet containing nuclei, was resuspended in nuclear extraction buffer (20 mM HEPES, pH 7.9; 420 mM NaCl; 1.5 mM MgCl_2_; 0.2 mM EDTA; 0.5 mM DTT; 0.5 mM PMSF) supplemented with protease inhibitors (Roche). This mixture was incubated at 4 °C and vortexed every 10 minutes over a total period of 50 minutes. Then it was centrifuged at 16,000 g for 20 minutes, and the supernatant was collected, aliquoted, snap-frozen and stored at −80 °C. Protein concentrations were measured using a BCA assay (ThermoFisher).

### DNA Probe Generation

#### PCR-based Asymmetric Probe Generation

VEGFA probe was amplified exactly as described in ref (21), hSP1 probe was amplified via PCR using Dreamtaq DNA Polymerase (Thermo Fisher Scientific) with biotinylated forward primer o6120, and reverse primer o6119. The asymmetrical probe generation and characterization was conducted as described in (21).

#### Synthetic Probe

mSP1 probe is generated by hybridizing 1.5 µM FAM and biotin labelled forward primer and 1.8 µM unlabeled reverse primer (**Table SI 1-2**) in IDT buffer (30 mM HEPES, 100 mM KOAc) boiling for 5 minutes and overnight cooling down to room temperature in a Dewar flask filled with boiled water.

### Enrichments

The enrichment experiments were conducted as described previously (21). Briefly, 5 µL of Dynabeads MyOne Streptavidin T1 (ThermoFisher Scientific, 65601) were incubated with 4 pmol of biotinylated DNA probe (and 15 µL beads and 12 pmol DNA probe for mSP1) in 1× B&W buffer for 3 h at room temperature. The beads were then washed once with 1× B&W buffer, followed by two washes with Protein Binding Buffer (PBB: 50 mM Tris-HCl, pH 8; 150 mM NaCl; 1 mM DTT; 1 mM PMSF; 0.25 % IGEPAL) supplemented with protease inhibitors. Subsequently, the beads were incubated with 50 or 100 µg of nuclear extracts (100 µg for HEK293T VEGFA enrichment experiment and 50 µg for rest) and 5 µg of dA:dT competitor DNA (1.1 µM) in PBB, in a total volume of 300 µL, on a turning wheel overnight at 4 °C. After incubation, the beads were washed twice with 1× PBS and stored in 20 µL PBS until further use. For proof-of-concept experiments, 0.1 nM spike-in proteins (MBP-MBD2 or MBP-MeCP2) were added alongside with the nuclear proteins,

### Western Blot

Pulldown samples were eluted in 15 µl PBS and boiled with 1x sample buffer at 95°C for 5 min and applied to 12 % SDS-PAGE. Then it is transferred to polyvinylidene fluoride (PVDF) membranes using a Trans-Blot Turbo system (BioRad, 1.9 A, 25 V, 7 min). Membranes were blocked for at least 1 hour at room temperature, using 5 % (m/v)-skimmed milk in TBST (TBS + 0.1 % v/v Tween-20). MBP-antibody (NEB #E8032) was incubated 1:10000, beta tubulin-antibody (Cell Signaling #2146) and lamin A/C (Cell Signalling #2032) antibody was incubated 1:1000 in 1.5 % skimmed milk in TBST overnight at 4°C. The secondary anti-mouse or rabbit antibody coupled to horseradish peroxidase (Sigma, GENA931 or GENA934 respectively) was incubated in 2 % skimmed milk in TBST for 1 hour at room temperature. Between each incubation, membranes were washed three times with TBST. The chemiluminescent reaction was established using a Clarity Western ECL substrate (Bio-Rad) and detected on a Bio-Rad ChemiDocTM imaging system.

### On-Bead Digestion and Desalting

The on-bead digestions and desalting are conducted exactly as described in(21). Pulldown samples on the beads stored in 20 µL PBS were denatured and reduced by adding 50 µL of denaturation/reducing buffer (8 M urea, 50 mM Tris, pH 7.5, 1 mM DTT) and incubating for 30 min at 20 °C with shaking at 350 rpm. Subsequently, 5.55 µL of alkylation solution (50 mM chloroacetamide in denaturation/reducing buffer) was added, and the mixture was incubated for another 30 min at 20 °C, 350 rpm. On-bead digestion was performed by adding 1 µg Lys-C (NEB) dissolved in nuclease-free water and incubating for 1 h at 37 °C with shaking at 350 rpm. The supernatant was collected into a fresh protein low binding tube (Eppendorf). Next, 1 µg Trypsin (NEB) in 165 µL 50 mM Tris-HCl was added to the beads and incubated for 1 h at 37 °C, 350 rpm. Supernatants from both digestions were combined, and an additional 2 µg Trypsin was added to the pooled sample for overnight digestion at 37 °C with shaking at 350 rpm. The reaction was stopped by adding 20 µL of 10 % (Trifluoroacetic acid) TFA. Peptides were desalted using Pierce C18 Tips (ThermoFisher). Tips were first activated by aspirating and dispensing 100 µL 50 % acetonitrile (ACN) in water twice, then equilibrated with 100 µL 0.1 % TFA. Samples were loaded by aspirating and dispensing 15 cycles, followed by rinsing with 0.1 % TFA/5 % ACN twice. Peptides were eluted by aspirating 20 µL of 0.1 % formic acid twice into a new tube. The eluted peptides were dried in a vacuum concentrator at 30 °C for 1.5 h.

### Mass Spectrometry Measurements

Each experiment comprised five conditions with five replicates each (n = 5), resulting in 25 samples per cell line–probe combination. In total, four cell line–probe combinations were analyzed. Samples from the same cell line and probe were processed and analyzed together, whereas different combinations were handled separately. Dried peptides were reconstituted in 20 µL of Buffer A (0.1% formic acid in water).

For each sample, 200 ng of peptides were loaded onto a Vanquish Neo UPLC system (Thermo Fisher Scientific) and separated using an 80 min gradient ranging from 5–60% acetonitrile in 0.1% formic acid. Eluting peptides were introduced directly via nano-electrospray ionization into an Orbitrap Exploris 480 mass spectrometer (Thermo Fisher Scientific, Waltham, MA, USA).

Data acquisition was performed in data-independent acquisition (DIA) mode, consisting of a full MS survey scan at 120,000 resolution, followed by 40 DIA segments (16.3 Th isolation width with 1 Th overlap) spanning a mass range of 350–1000 m/z. MS/MS spectra were acquired at 30,000 resolution, resulting in a cycle time of 3 s and an average of ∼6 data points per chromatographic peak. The automatic gain control (AGC) target was set to 300% for MS and 1000% for MS/MS scans, with a maximum injection time of 45 ms for MS scans. Fragmentation was performed using higher-energy collisional dissociation (HCD) at a normalized collision energy of 30%.

### Mass Spectrometry Data Analysis

Raw data were processed using DIA-NN (version 1.8.2) (23) with a UniProt reference proteome database for *Homo sapiens* (HEK293T VEGFA and hSP1 probe; HeLa VEGFA probe) or *Mus musculus* (mouse embryonic stem cells with mSP1 probes) (release June 2024). Oxidation (M), protein N-terminal acetylation, and methionine excision were set as variable modifications, while carbamidomethylation (C) was defined as a fixed modification. Up to one missed cleavage was allowed, with a maximum of three variable modifications per peptide. False discovery rates (FDR) were controlled at 1% at both peptide and protein group levels. The resulting report.pg_matrix.tsv output was further analyzed using Perseus (version 2.0.3.1) (24). Protein intensities were log_2_-transformed, and samples were grouped according to experimental condition. To ensure data integrity, samples that did not meet predefined quality criteria were excluded from the analysis (excluded samples were for the HEK293T-VEGFA experiment: mC/mC replicate 5 (sample 16) and fC/mC replicate 1 (samples 17), for the HEK293T-hSP1 experiment: C/C replicate 3 (sample 4), and for the mESC-mSP1 experiment: C/C replicate 5 (sample 6) and fC/fC replicate 5 (sample 26)); the number of replicates may therefore vary across samples but a minimum of four replicates was retained for all conditions.

For pairwise comparisons, proteins were required to have more than four valid values in at least one group. Missing values were subsequently imputed from a normal distribution (downshift 1.8, width 0.3). Statistical significance was assessed using a two-sided Student’s t-test with permutation-based multiple testing correction. Proteins with a log_2_ fold change > 2 and an adjusted p-value < 0.01 were considered significantly enriched. Volcano plots were generated using VolcaNoseR (25).

For statistical analysis of the complete dataset (5 conditions for one probe), proteins were retained only if at least four replicates per condition afforded a valid LFQ value. An ANOVA multiple-sample test with a permutation-based FDR of 0.01 was performed to identify significantly enriched proteins. Missing values were imputed from a normal distribution (downshift 1.8, width 0.3). The corresponding LFQ values were z-scored and visualized in a heat map.

### Cloning and Purification of Recombinant Proteins

#### Cloning

The plasmids of MAX and RFX5 DBD are used from(21). The coding sequences of HEY1 (aa2-304, [NM_012258.4]), SIX2 (aa2-291, [NM_016932.5]), SIX1 (aa2-284, [X91868.1]), FOXJ3 (aa2-588, [NM_001198852.2]), TP53 (aa2-393, [NM_001126112.3]), MPG (aa1-298, [NM_002434.4]), TDG (aa1-410, [NM_003211.6]), APEX1 (aa2-318, [NM_001641.4]), and XRCC1 (aa1-633, [NM_006297.3]) are amplified from thyroid (HD-503, Zyagen) or prostate (10108-A, Biocat) cDNA libraries (see **Table SI 3** for primer sequences) and cloned into an XhoI-digested pET-21d(+) vector with N-terminal MBP tag and C-terminal His tag by Gibson assembly.

#### Purification

Expression plasmids were transformed into *E. coli* BL21-Gold(DE3) by heat-shock (Agilent). Single colonies were selected from agar plates with LB-Miller broth supplemented with 50 µg/mL carbenicillin (LB Carb) and inoculated into LB Carb liquid medium for overnight pre-growth. Then the culture was diluted to an optical density (OD600) of 0.05 in 30 mL LB Carb medium and grown at 37°C until reaching to an OD600 of 0.6–0.7. Expression was induced with 1 mM IPTG and supplemented with 1 mM MgCl_2_, and 1 mM ZnSO_4_, and cultures were incubated overnight at 25°C or 30°C, shaking at 150 rpm. Cells were harvested and washed once with ice-cold 20 mM Tris-HCl (pH 8.0) and resuspended in 2 mL binding buffer (20 mM Tris-HCl pH 8.0, 250 mM NaCl, 10% glycerol, 10 mM 2-mercaptoethanol, 5 mM imidazole, and 0.1% Triton X-100) with 1 mM PMSF. Resuspended cells were sonicated (Branson Digital Sonifier 450 Cell Disruptor, 3 min, amplitude 15 %, 4 s on, 2 s off) and subsequently incubated with 1 mg/mL lysozyme and 10 U DNAseI (NEB M0303S) overnight at 4°C. After centrifugation at 14000 × g for 20 min at 4°C, the supernatants were collected and incubated at 4°C for 20 min, shaking at 700 rpm with 450 µL 50% Ni-nitriloacetic acid (NTA) HisPur agarose resin (ThermoFisher) that was previously equilibrated with binding buffer. The resins were washed 2 × with 1 mL binding buffer containing 90 mM imidazole for 5 min, and protein was eluted in 2 × 0.2 mL and 1 × 0.4 mL binding buffer with 500 mM imidazole for 5 min. The purify was evaluated by running SDS PAGE (**Figure SI 25**), and successful elutes were combined and dialyzed with dialysis buffer 3 times (20 mM HEPES pH 7.3, 100 mm NaCl, 10% glycerol, and 0.1% Triton X-100 in a tube with 12–14 kDa MWCO (Roth). The protein concentration was determined with a BCA assay (ThermoFisher) and the proteins were snap-frozen in liquid nitrogen and stored at −80°C.

### Electrophoretic Mobility Shift Assays

VEGFA and mSP1 DNA probes with FAM labels are obtained as described *in DNA probe generation* section. MBP-fusion proteins were treated with 0.25 µm TEV-protease overnight at 4°C to cleave off the MBP tag. For protein binding studies, various concentration of proteins was incubated with 2 nM labeled DNA probe with indicated modifications in EMSA buffer (20 mM HEPES pH 7.3, 30 mM KCl, 1 mM EDTA, 1 mM (NH_4_)_2_SO_4_, 0.2% Tween-20) in presence of 5 µM dA:dT competitor and 1 mM DTT for 20 min at 21°C. For titration experiments, a dilution series of protein was prepared, ranging from 0.5 - 512 nM for MAX, 0.5 – 512 nM for HEY1, 4 - 4096 nM for SIX2, 4 - 4096 nM for SIX1, 2-2048 nM for FOXJ3, 4 - 4096 nM for TP53, 0.5 – 512 nM for MPG, 0.5 – 256 nM for TDG, 2-2048 nM for APEX1, and 2-2048 nM for XRCC1. Proteins were incubated with 2 nM labeled DNA probe as described above. Then samples were mixed with EMSA loading buffer (1.5X TBE with 40% glycerol) and loaded to non-denaturing polyacrylamide gel (12% for VEGFA probe, 15% for mSP1 probe) that was pre-run at 220 V for overnight at 4°C. After loading the samples, gels were run at 240 V at 4°C for 75 min, and visualized by the 510 LP filter of the Typhoon FLA-9500 laser scanner (GE Healthcare) at 473 nm with 800 V-1000 V amplification. Bound and unbound DNA probes were quantified using ImageQuant TL v8.1 1D Gel Analysis (GE Healthcare). The bound fraction of DNA was calculated for each protein concentration and analyzed as described in ref (21).

### Modelling

For modelling of the protein-DNA interactions, AlphaFold 3 was used to predict the structure with protein sequences given above and mSP1 probe by positioning the non-canonical cytosines accordingly (26). Figures were composed with Pymol 2.5.4 (27).

## RESULTS

### Enrichment/MS-based proteomics for discovering (anti)readers of symmetric and asymmetric fC-modifications

To obtain proteome-wide interaction profiles of fC-modified CpG dyads, we used an enrichment protocol coupled with quantitative HPLC-MS/MS measurements. We employed a ∼200 nt long enrichment DNA probe containing part of the promoter region of human vascular endothelial growth factor (VEGF) A gene covering a high density of TF binding sites and 11 CpGs within different sequence contexts (**Figure 2a**) (21). VEGFA is a pivotal regulator of angiogenesis (28), its expression is controlled by promoter methylation and TET3 (29), and aberrant methylation is associated with cancer and inflammation (30-32). We generated five different versions of the probes being strand-symmetrically or - asymmetrically modified with C, mC or fC, resulting in probes containing the CpG dyads C/C, mC/mC, fC/C, fC/mC and fC/fC. All probes were PCR-generated with a biotinylated primer as described previously using dCTP, dmCTP or dfCPT in the dNTP mix (see ref (21) for a discussion of probe design parameters). Nuclear proteomes were extracted from HEK293T cells (**Figure SI 1a-d**) and incubated with DNA probes immobilized on streptavidin beads. After washing, bound fractions were eluted and analyzed either by western blot in proof-of-concept experiments or by HPLC-MS/MS for identifying proteome-wide reader profiles (**Figure SI 2a**). For the proof of concept, we added a small amount of a fusion protein of maltose binding protein and the well-studied mC/mC reader protein methyl-CpG-binding domain protein 2 (MBP-MBD2) as a spike-in to a nuclear proteome and analyzed the input and enriched fractions by anti-MBP western blots. MBD2 was most highly enriched by the symmetric mC/mC probe followed by the asymmetric mC/fC probe, whereas other probes showed only little or no enrichment, faithfully resembling MBD2’
ss CpG dyad state preferences previously observed in electromobility shift assays (EMSA) (**Figure 2b** and **SI 2b,d**) (33). The same experiment conducted with a second MBD-based mC/mC reader (MBP-MeCP2) revealed a sightly distinct profile than obtained for MBD2, again being in close agreement with previous EMSA data (33) (**Figure SI 2c,e**). These results indicate the capability of our enrichment protocol to reveal even subtle differences in reader’s CpG dyad preferences. Next, we enriched and analyzed nuclear proteomes with HPLC-MS/MS to identify their binding preferences (see **Figure SI 3** for quality control data). Overall, the highest number of significantly enriched proteins was detected for the fC/C probe followed by the symmetric C/C, mC/mC and fC/fC probes, whereas the lowest number was observed for the fC/mC probe (**Figure 2c**).

**Figure 2.**
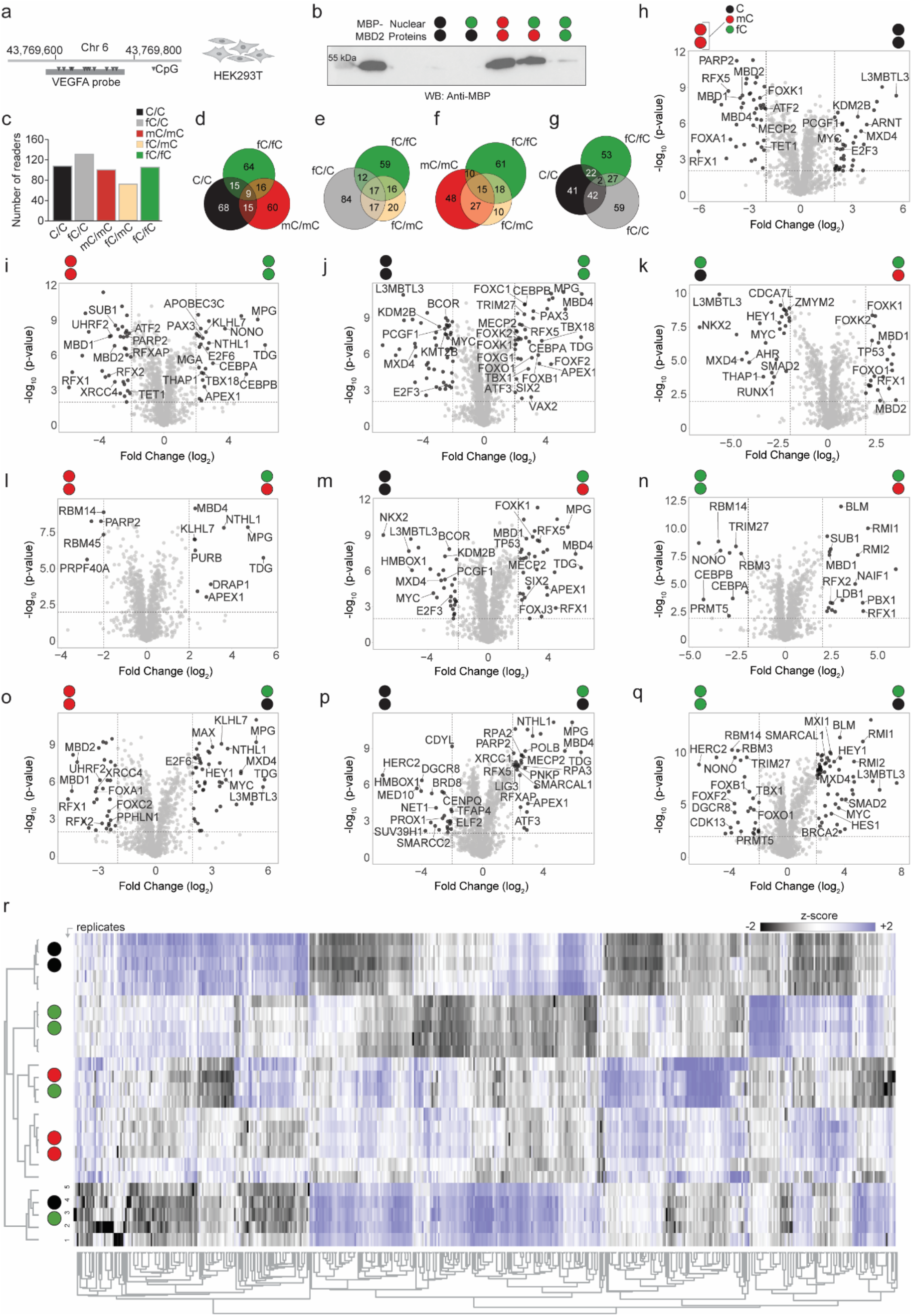
Discovery of human readers and antireaders of symmetric and asymmetric fC-modifications by proteomics. (a) Employed probe design (**VEGFA**) and nuclear extract **(HEK293T)** used in this experiment. (b) Anti-MBP western blot of the enrichment of MBP-MBD2 spike-in from nuclear lysate using VEGFA probes modified as indicated. (c) Overall number of significantly enriched reader proteins for indicated probe modifications. (d-g) Venn diagrams showing the overlaps of significantly enriched proteins. Note that **Figure 2c-g** use a color code describing both DNA strands at once. (h-q) Volcano plots comparing pairwise binding partners of the indicated probes. The -log10 p-value (Student’s t-test; y-axis) is plotted against the log2 fold change (x-axis). Threshold for significance: -log10 p-value < 0.01, log2 fold change ≤ -2 or ≥ 2 (see Table SI 7 for details). (r) Proteins were retained only if at least 4 replicates per condition afforded a valid LFQ value. ANOVA (permutation-based FDR 0.01) was performed, and significant proteins were z-scored and visualized in a heat map. Missing values were imputed from a normal distribution (downshift 1.8, width 0.3). For cytosine modification colour code, see **Figure 1**.

Whereas a high number of fC readers was also observed in previous studies limited to symmetric fC/fC probes (16-18), our new data with fC/C and fC/mC probes indicates that fC also attracts many proteins in asymmetric contexts. Next, since proteins may have the ability to bind to more than one probe, we analyzed the reader overlap between probes. Strikingly, although some overlap was observed in all cases, the five probes enriched rather distinct reader groups (**Figure 2d-g**). We next made pairwise comparisons of enrichments with individual probes by generating volcano plots (**Figure 2h-q**). As an initial validation of our method, we first compared enrichments with C/C versus mC/mC probes, since the readers of C and mC nucleobases are well characterized and several reference studies are available (16-18,21,34) (**Figure 2h**; see also **Table SI 7** for comparisons from **Figure 2h-q** showing all protein names). In agreement with the literature, MBD proteins (MBD1,2,4 and MeCP2), RFX family proteins (RFX1,2,5,7, RFXAP and RFXANK), FOX family proteins (FOXA1, FOXB1, FOXC1-2, FOXF2 and FOXK1-2), ATF proteins (ATF2,7), UHRF2 and TET1 were enriched for mC/mC. Moreover, the C/C probe enriched readers previously found for unmodified DNA probes, such as subunits of the polycomb repressive complex (PRC; e.g., PCGF1, BCOR and KDM2B), as well as its functional cooperation proteins (L3MBTL3, CDYL, MXI1 and MXD4) (35) (**Figure 2h**). This data confirms that our probes and enrichment assay work as expected on the proteome level.

Next, we compared fC/fC and the two other symmetric probes (**Figure 2i-j**). For mC/mC and fC/fC probes, the former enriched typical mC readers, whereas mainly DNA repair proteins (TDG, MPG, APEX1, NTHL1, APOBEC3C) and TFs (CEBPA,B, TBX18, PAX3, E2F6, MGA and THAP1) were enriched for the fC/fC probe (**Figure 2i**). Similarly, when compared to the fC/fC probe, the C/C probe showed largely an enrichment of the same C readers observed in the C/C versus mC/mC probe comparison (**Figure 2j**). However, the fC/fC probe did not only enrich DNA repair proteins and TFs (ATF3, CEBPA,B, PAX3, SIX2, TBX1,18, VAX2, FOXG1, FOXO1), but also proteins that were enriched by the mC/mC probe before, indicating a certain tolerance of these mC readers for fC. This included MBD proteins (MBD4 and MeCP2), RFX5 proteins (RFX5) and several FOX proteins (FOXB1, FOXC1, FOXF2, FOXK1-2) (compare **Figure 2h, j**). We next compared fC/mC probes with all four other probes (**Figure 2k-n**). Here, readers of fC/C probes were higher in number than readers of fC/mC probes (**Figure 2c**), and interestingly, we also observed much higher fold changes for the former, illustrating a strong preference of many proteins for fC/C over fC/mC probes (**Figure 2k**). Significantly enriched proteins for the fC/C probe covered typical C readers (L3MBTL3, MYC and MXD4, compare with **Figure 2h**), but also showed several unique readers, such as CDCA7L, ZMYM2 and HEY1. In contrast, the proteins enriched by the fC/mC probe mainly covered typical mC/mC readers, such as MBD1-2, RFX1, and FOX family proteins, again indicating a certain tolerance for fC. Interestingly, the tumor suppressor TP53 showed a clear symmetry preference by being enriched for fC/mC over fC/C (**Figure 2k**) and also C/C probes (**Figure 2m**). In a comparison between asymmetric fC/mC and symmetric mC/mC probes, the latter enriched only few proteins such as RBM14,45 and PARP2, whereas the fC/mC probe enriched BER proteins (TDG, MPG, NTHL1, APEX1) which were overlapping with the aforementioned fC/fC reader proteins (**Figure 2l**). This included the repair glycosylase MBD4, whereas no other MBD was enriched for either dyad, being in agreement with a relatively high affinity of many MBDs for fC/mC dyads (33). Strong differences were observed between the fC/mC and C/C probes. The latter showed several C readers observed also in comparisons with mC/mC probes, whereas the former enriched a mix of repair proteins (**Figure 2m**; MPG, MBD4, APEX1, TDG) and mC readers (RFX1,2, FOXK1, FOXJ3, MECP2). Comparing fC/mC with symmetric fC/fC probes, the latter generally enriched a higher number of readers (**Figure 2c**). Enriched readers included CEBPA,B, RBM3,14 and NONO, whereas the fC/mC probe enriched the BTR complex members BLM and RMI1,2 (36), MBD1, RFX1,5 and SUB1 (**Figure 2n**). Finally, we made comparisons between the asymmetric fC/C probe and the remaining probes (**Figure 2o-q**). The comparison between the fC/C and mC/mC probe showed similarities with the previous fC/C versus fC/mC comparison (compare **Figure 2o** and **k**, overlapping proteins for fC/C in both cases included MYC, MXD4, L3MBTL3 and HEY1). However, in the new comparison, fC/C showed enrichment of KLHL7, E2F6 and MAX along with repair proteins (TDG, MPG and NTHL1), presumably given the absence of an fC in the compared probe. Compared to the C/C probe, the fC/C probe enriched many DNA repair proteins including most components of the BER complex (37) (TDG, APEX1, LIG3, XRCC1, POLB, PARP2, PNKP, MPG, NTHL1, RPA2,3 and SMARCAL1), but also MBDs (MBD4 and MeCP2) and RFX family proteins (RFX5 and RFXAP) (**Figure 2p**). In contrast, the C/C probe enriched TFs (HMBOX1, PROX1, ELF2 and TFAP4) and other chromatin regulators (BRD8, CDYL, SMARCC2 and SUV39H1). Finally, when comparing the fC/C and fC/fC probes, the asymmetric fC/C probe showed higher fold changes (**Figure 2q**) and an overall higher number of readers (**Figure 2c**). fC/fC enriched several TFs (CEBPA,B, FOXB1, FOXF2, FOXO1 and TBX1) and RBM3,14), whereas fC/C enriched DNA repair proteins (BRCA2, BLM, RMI1,2 and SMARCL1), leucine zipper-basic helix-loop-helix (LZ-bHLH) TFs (MYC, MXD4, HES1 and HEY1) and L3MBTL3. Members of the BTR complex (BLM and RMI1,2) were also enriched for mC/fC when compared to fC/fC (36) (**Figure 2n**).

In a hierarchically clustered heatmap of the combined enrichment data, distinct modification preferences of many proteins and several general trends can be observed (**Figure 2r**; shown are significantly enriched proteins from ANOVA test). For example, though showing individual protein groups with specific preferences between the two probes, fC/mC and mC/mC data overall cluster closely together, suggesting that fC and mC are recognized with comparably similar affinities by many proteins. Indeed, whereas the fC/fC data is already more distant from the mC/mC data, fC/mC and fC/fC also cluster closely (**Figure 2r**). In contrast, C/C and mC/mC data are segregated in distinct clusters, as somewhat expected from their overall opposing regulatory functions and in agreement with previous data (21). Remarkably, the rather similar C/C and fC/C probes resulted in the most distinct segregation, with many proteins showing very pronounced enrichment differences (**Figure 2r**). Finally, for obtaining information on how readers are functionally connected, if they may be enriched directly or indirectly as part of complexes, and to what class of proteins they belong, we conducted STRING protein–protein interaction network analyses for the readers of each probe (38,39). Readers of C/C and mC/mC probes were mainly TFs and chromatin regulators, but no or little BER proteins, respectively (**Figure SI 4** and **6**, respectively). All fC probes attracted prototypical BER proteins (TDG, APEX1, POLB, MPG etc.) along with TFs and chromatin regulators (**Figure SI 5,7,8**). Interestingly, the fC/fC probe readers contained a wide range of loosely connected TFs including a cluster of TFIID members involved in transcription initiation (**Figure SI 8**).

### Sequence dependence of (anti)readers of fC-modifications

To gain insights into potential DNA sequence-dependencies of fC (anti)readers, we conducted comparative enrichments with the same HEK293T proteomes, but with a different enrichment probe design. The new probe contained a promoter region of the human specific protein 1 (hSP1) gene and was similar in size and in the number of contained CpG dyads as the VEGFA probe (**Figure 3a, Table SI 6** shows JASPAR analysis of TF sites). SP1 is a TF that is highly upregulated during early embryogenesis (40) and it regulates the expression of multiple genes including itself. It is a central target for proliferation and embryonic development (41) and is tightly regulated during cell cycle and upregulated by overexpression of TET1 (41-43). We generated the five different probe versions as before and again confirmed their abilities to faithfully report the modification preferences of MBD2 by spike-in enrichments and western blots (**Figure SI 10b**) (33). We then conducted and analyzed enrichment experiments by HPLC-MS/MS as before (see **Figure SI 9** for quality control data). Strikingly, the overall distribution of reader numbers enriched for each group was very similar as observed for the VEGFA probe, with similar overlaps between modifications (compare **Figure SI 10c-g** and **Figure 2c-g**). However, though volcano plots revealed a consistent divergence pattern as seen before, as well as a larger number of proteins that were enriched for the same modifications in both probe designs, there was also a larger number of new proteins not enriched with the VEGFA probe. For example, SIX1,2, SATB1,2, CDCA7L and CDCA7, DDX1, TFDP1, PAX9,2, VAX1, RTRAF, KDM2B, RFC3 and ING5 all showed an fC-dependent enrichment with varying preferences for specific fC modification symmetries of the probes (**Figure SI 10i-q** and **Table SI 8**). The hierarchically clustered heat map again showed unique probe signatures of many proteins (**Figure SI 10r**). In conclusion, we observe similar distributions of reader numbers and similar reader overlap patterns for the two probe designs. Whereas some proteins are enriched for both probes and may exhibit sequence-independent read-out of the modifications, there is also a large number of probe/sequence-specific readers (**Figure 3b** shows the Venn diagrams of the probe comparison, **Table SI 4** shows the protein lists). This indicates that many more sequence-specific fC readers may exist and could be discovered via additional probe designs.

**Figure 3.**
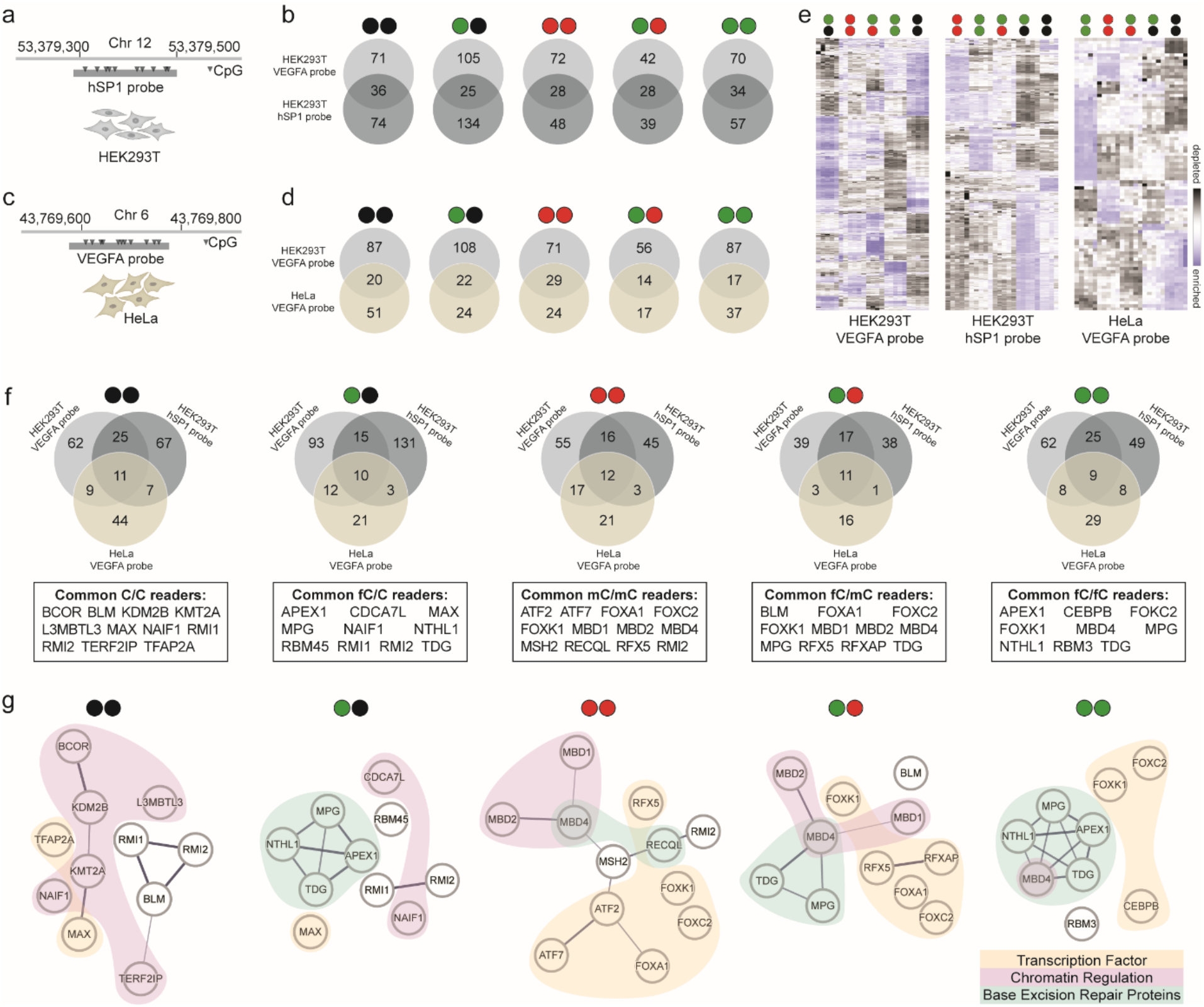
Overview and common (anti)readers from combined human proteomics experiments. (a-b) Experiment setups with different probe designs and corresponding Venn diagrams showing the overlap of readers between setups. (c-d) Experiment setups with different cell lines and corresponding Venn diagrams showing the overlap of readers between setups. (e) Comparison of heatmaps from all human experiments, for detailed description see **Figure 2r** and **Figure SI 10r** and **12r**. (f) Venn diagrams showing the overlap between human experiments with lists of common readers below. (g) Interaction network of common reader groups with colour code shown on the bottom left. Analysed with STRING v12.0. For cytosine modification colour code, see **Figure 1**.

### Tissue dependence of (anti)readers of fC-modifications

To discover additional readers and to evaluate potential tissue-dependence of fC preferences of already identified readers, we next conducted enrichments with nuclear extracts of HeLa (human cervical adenocarcinoma) cells employing our VEGFA probe (**Figure SI 1e-g** and **12b;** see **Figure SI 11** for quality control data). We observed a similar distribution of reader numbers for fC-modified probes and similar reader overlaps as in previous experiments, with however inverted numbers for C/C and fC/C probes (**Figure SI 12c–g**). Volcano plot analyses revealed subsets of proteins exhibiting unique enrichment profiles in response to different CpG modifications (**Figure SI 12h–q** and **Table SI 9**). We observed a number of proteins showing similar probe preferences as observed for HEK293T proteomes. However, a larger number of proteins was HeLa-specific, with, e.g., ORC1, WDR5, CBX5, PURA,B, ARNT, AHRR, NAIF1, several HNRNPs and many other showing a preference for individual or multiple fC-modified probes (**Figure 3d** and **Table SI 5** shows the Venn diagrams and protein lists for this experiment). Hierarchical clustering further delineated distinctive binding signatures (**Figure SI 12r**). These findings indicate a substantial tissue-dependence of reader enrichment, in agreement with studies employing symmetric fC or other oxi-mC modified probes. This again suggests that many more fC readers may exist and could be discovered by studying additional cell lines/tissues.

### Overview of (a)symmetric fC (anti)readers in the human proteome

In our profiling experiments with human proteomes employing two distinct DNA probe sequences and nuclear extracts from two cell lines, we observed a pronounced sequence- and tissue-dependence of reader profiles (**Figure 3e** shows a comparative overview of the heatmaps). Despite these variations, we observed a small subset of proteins for each probe modification type that was consistently enriched across all experiments with comparable abundance ranges. These were designated as common readers for the respective probes (**Figure 3f** shows Venn diagrams and reader lists). Although the (a)symmetric dyads fC/C and fC/mC shared partial overlap with the C/C and mC/mC probes, respectively, they also attracted many unique readers (**Figure 3f**) and exhibited differences in enrichment magnitudes (**Figure 2** and **SI 10,12)**. To study potential broader functions of different fC modifications, we explored the functional relationships among the common readers by STRING analyses (**Figure 3g**) as conducted before. The common C/C readers predominantly comprised chromatin regulatory proteins, whereas common fC/C readers were enriched in BER factors. In contrast, common mC/mC and fC/mC readers displayed more heterogeneous functional distributions with a subtle enrichment for TFs. Notably, common fC/fC readers exhibited a marked reduction in chromatin regulators but included numerous BER proteins and TFs. We further analyzed the networks for GO enrichment via STRING (38,44,45). The results depict how readers are involved in general biological processes and the degree of significance (**Figure SI 13-17**; for complete GO terms, see **Table SI 10**). Distinct GO term clusters emerged for each modification type, suggesting specific functional roles.

### Discovery of the (anti)readers of symmetric and asymmetric fC modifications in mESC

In mouse early embryogenesis, fC accumulates in the paternal pronuclei shortly after fertilization (46) and occurs as intermediate of an active demethylation of the paternal genome, but can also occur as a stable mark (47). mESC exhibit high fC levels (48), but have so far been profiled only with symmetric fC probes. We designed a new DNA probe covering part of the promoter region of the mouse Sp1 gene (designated mSP1). We kept this probe relatively short (50 nt including four CpG sites), to enable its generation through solid phase DNA synthesis (**Figure 4a**). Being four times shorter and containing a lower number of protein binding sites than our previously used probes, this synthetic probe is expected to enrich correspondingly fewer proteins (see **Table SI 6** for JASPAR analysis of TF sites). However, in contrast to the standard PCR-based approach for DNA probe generation (17,18,21), solid phase synthesis (though extremely costly for fC probes) enables C modifications to be selectively incorporated only at CpG sites. This DNA probe is the first one in the oxi-mC proteomics field that features a complex, natural sequence and contains modifications only at CpGs (the only other synthetic DNA probe previously employed in the field is a 27mer containing four C modifications in an (ACG)_4_ repeat sequence) (16-18). We next extracted nuclear proteomes from mESCs (**Figure SI 1h–j**) and conducted enrichments with the mSP1 probe as before (**Figure 4b** shows MBP-MBD2 control; see **Figure SI 18** for quality control data). We observed the highest number of enriched readers for the fC/fC probes followed by mC/mC and C/C. fC/mC and fC/C probes enriched less proteins, in combination nevertheless exceeding the number of the canonical C/C or mC/mC probes, respectively (**Figure 4c**). The Venn diagrams showed a comparable overall picture as observed for the human enrichments, with relatively little overlap between probes (highest overlaps were observed between fC/mC and mC/mC or fC/fC probes, respectively) (**Figure 4d–g**). Pairwise comparisons by volcano plots illustrated that a single change in the CpG strand modification resulted in significantly altered reader profiles in all cases (**Figure 4h–q**). Comparisons between mC/mC and C/C probes revealed enrichment of MBD proteins (Mbd1,2,4 and Mecp2), Uhrf1 and Clk2 for mC/mC and PRC subunits (Pcgf1, Bcor and Kdm2b), associated protein L3mbtl3 and antagonist Cxxc1 alongside with other proteins for C/C dyads (**Figure 4h;** see **Table SI 11** for protein lists of all comparisons). Comparing these two symmetric probes with symmetric fC/fC probes showed similar proteins enriched for mC/mC and C/C as in their direct comparison. However, BER proteins (Tdg and Apex), FOX family proteins (Foxp1,2,4 and Foxf2), RNA binding proteins (Rbms1,2, Rbm3, Pcbp1, Dazap1 and Lin28b) and others were enriched for fC/fC (**Figure 4i-j**). Interestingly, in contrast to the human experiments, fC/mC enriched a larger number of proteins than the other asymmetric probe fC/C (**Figure 4k**). Atf7 and Cdca7l were enriched for fC/C, whereas MBD proteins (Mbd1,2,4), FOX family proteins (Foxp1 and Foxk1,2) and Chaf1a were enriched for fC/mC over fC/C. Comparisons between mC/mC and fC/mC probes showed enrichment of Clk2 for the former and of BER proteins (Tdg and Apex) and Lin28b for the latter (**Figure 4l**). Compared to C/C and fC/fC, the fC/mC probe enriched typical mC readers and other new proteins (Mbd2, Mbd4, Uhrf1, Parp2, Lin28b, Topbp1 and Mbd1, respectively, **Figure 4m,n**). In comparisons of fC/C with other probes, probes containing C/C and mC/mC dyads enriched similar proteins as in their direct comparisons, whereas the fC/C dyad probes enriched repair proteins and Mxi1 as well as Topbp1 and Tp53, respectively (**Figure 4o-p**). We obtained a very different picture in the comparison with fC/fC dyad probes – here, many Fox proteins along with Ptb2 and Hdac3 were enriched for the symmetric dyad, whereas Atf7, Creb5 and L3mbtl3 were enriched for the fC/C dyad (**Figure 4q**).

**Figure 4.**
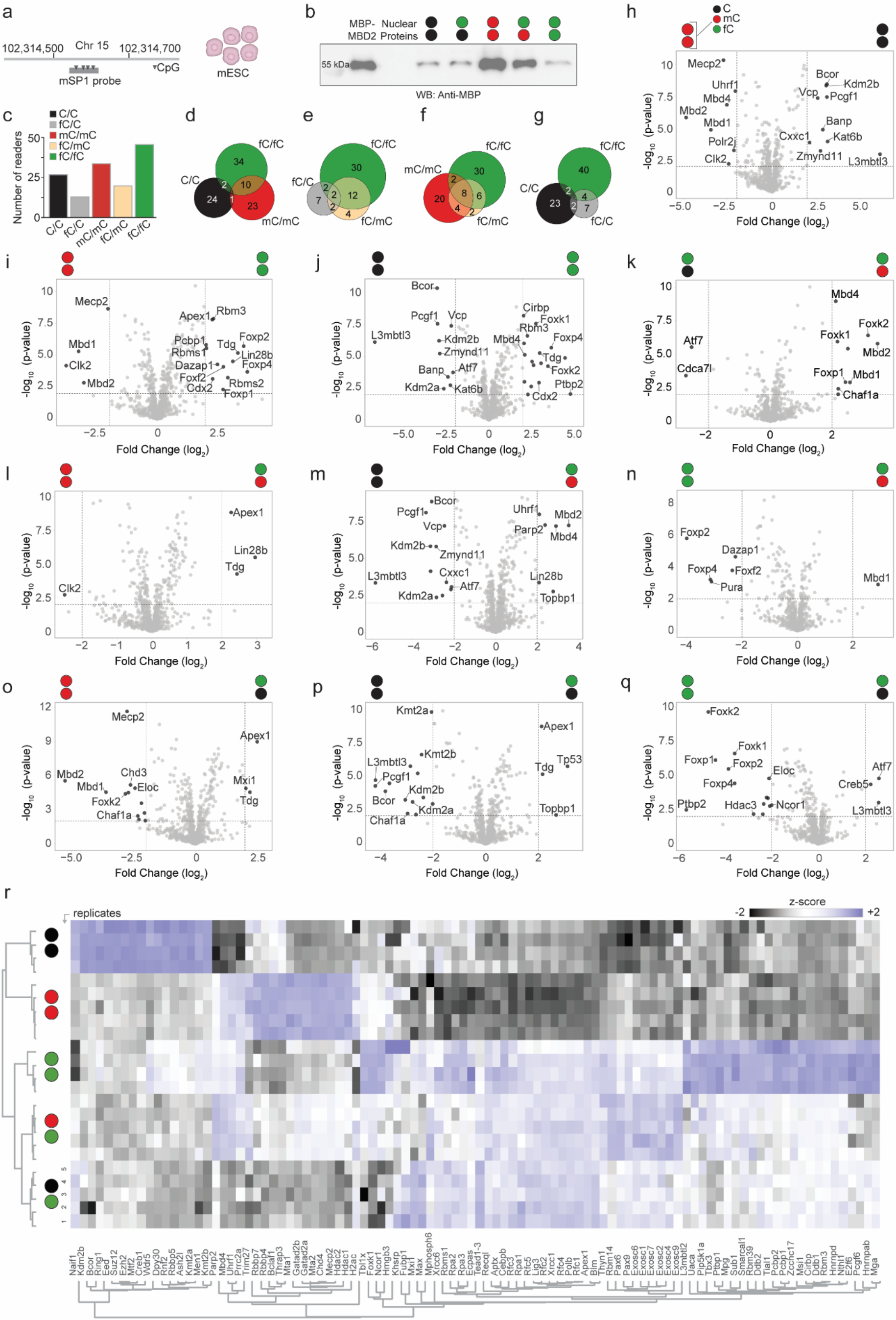
Discovery of mouse readers and antireaders of symmetric and asymmetric fC-modifications by proteomics. (a) Employed probe design (**mSp1**) and nuclear extract **(mESC)** used in this experiment. (b) Anti-MBP western blot of the enrichment of MBD2-MBP spike-in from mESC nuclear lysate using mSp1 probes modified as indicated. (c) Overall number of significantly enriched reader proteins for indicated probe modifications. (d-g) Venn diagrams showing the overlaps of significantly enriched proteins. Note that **Figure 2c-g** use a colour code describing both DNA strands. (h-q) Volcano plots from pairwise comparisons (log2-fold changes > 2 and p-value < 0.01; see **Table SI 11** for details). (r) Proteins were retained only if at least 4 replicates per condition afforded a valid LFQ value. ANOVA (permutation-based FDR 0.01) was performed, and significant proteins were z-scored and visualized in a heat map. Missing values were imputed from a normal distribution (downshift 1.8, width 0.3). For cytosine modification colour code, see **Figure 1**.

The hierarchically clustered heat map again demonstrated distinct binding profiles of proteins (**Figure 4r**). To understand the functional connections between the readers of each CpG dyad modification, we again conducted STRING analyses (38,39) (**Figure SI 19**). The protein network of C/C and mC/mC CpG dyad readers contained mostly chromatin regulators and several TFs (**Figure SI 19a,c**). As expected, BER proteins appeared at highest fractions for fC-modified CpG dyads. However, many TFs and to a smaller extent also chromatin regulators were enriched (**Figure SI 19b,d,e**), with the highest fraction observed for fC/fC CpG dyads (**Figure SI 19e**). We further analyzed the networks for GO enrichment via STRING (38,44,45), and again observed distinct GO term clusters for each probe type (**Figure SI 20-24**; for complete GO terms, see **Table SI 12**).

### Validation of (anti)readers

Our proteomics data afforded many proteins that were enriched with specific fC modification preferences. A number of proteins and protein classes have been observed in previous experiments using symmetric fC/fC probes as well as different probe sequence contexts and partially also cell lines/tissues. These included the repair proteins MPG, TDG, but also TFs like p53 and different FOX proteins (16-18,21). Interestingly, our new proteomics data suggest that several of these proteins show preferences for particular fC-modification symmetries of probes. Moreover, we observed many proteins with specific fC probe preferences that were not identified before. To evaluate if these (anti)readers were enriched via direct probe interactions or indirectly as part of larger complexes and to characterize their modification preferences in detail, we selected eleven reader candidates for binding analyses. These included TFs (**Figure 5)**, and BER proteins (**Figure 6**; see also **Figure SI 27**). We cloned and expressed the proteins as fusion proteins with an N-terminal MBP and a C-terminal His6 tag in *E. coli*, purified them (**Figure SI 25**), and conducted EMSA with the VEGFA and/or mSP1 enrichment probes. It should be noted that DNA-binding properties of a protein may depend on post-translational modifications (PTMs), other interacting proteins, RNAs, or ligands that are present only in mammalian nuclear lysates. Consequently, assays using recombinant proteins complement the proteomics data, but do not necessarily rule out candidate readers when negative.

**Figure 5.**
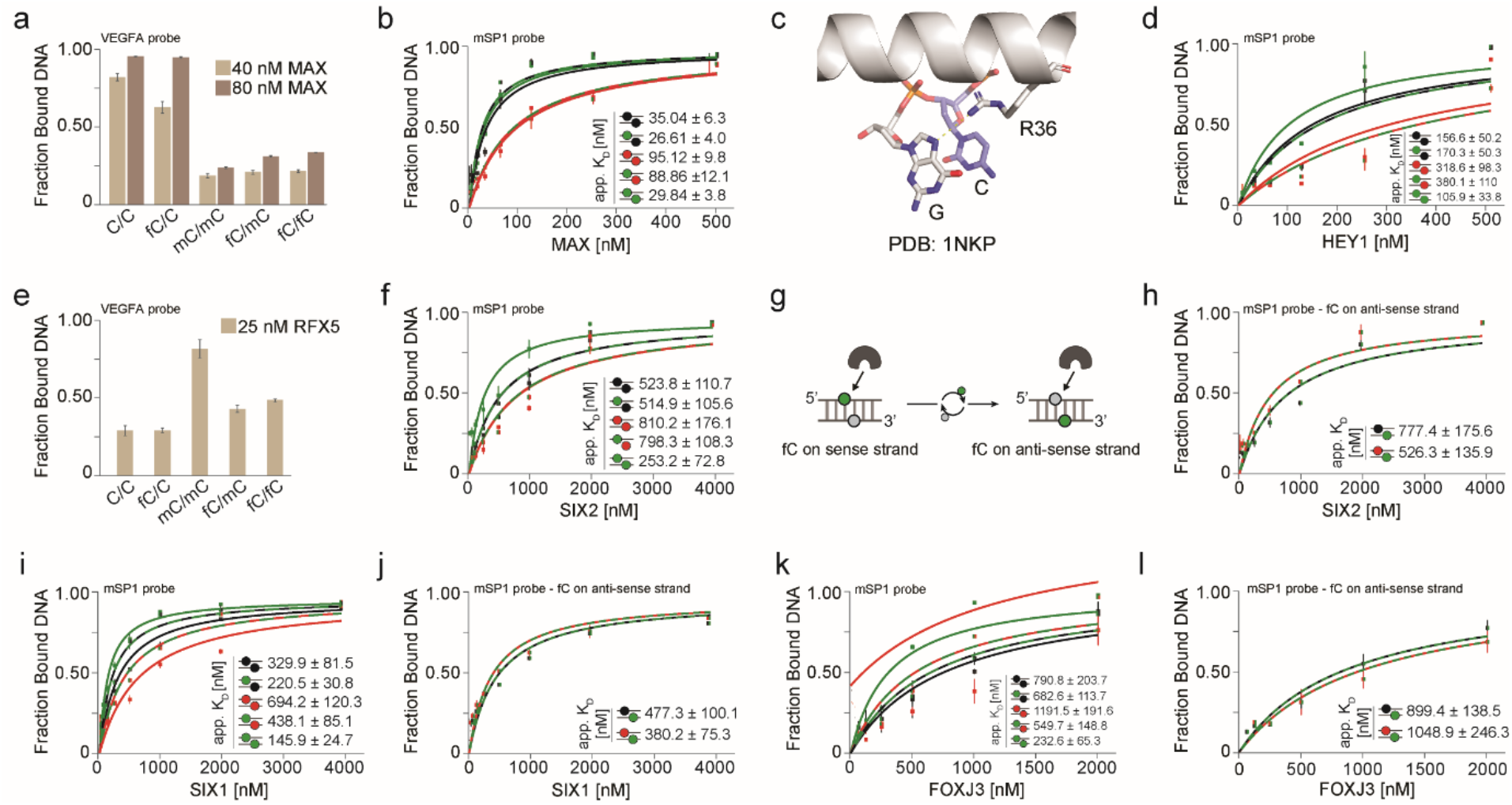
Validation of TFs reading fC-modified CpG dyads with distinct symmetry preferences. (a) EMSA profile for MAX_2_ binding to VEGFA probes. (b) EMSA titration experiment with MAX_2_ and mSp1 probes. (c) Crystal structure of MYC/MAX (pdb: 1NKP) bound to DNA showing conserved arginine (R36 in MAX) involved in binding the CpG dyad. EMSA profile for RFX5-DBD binding to VEGFA probes. (d) EMSA titration experiment with HEY1 binding to mSp1 probes. (e) EMSA profile for RFX5 binding to mSp1 probes. (f) EMSA titration experiments for SIX2 binding to mSp1 probes. (g) Cartoon illustrating strand swapping experiment. (h) EMSA titration experiments for SIX2 binding to mSp1 probes with fC on the antisense strand. (i-j) EMSA titration experiments for SIX1 binding to mSp1 probes, with fC on the sense (i) or antisense (j) strands. (k-l) EMSA titration experiments for FOXJ3 binding to mSp1 probe, with fC on the sense (k) or antisense (l) strands. Each experiment was done in duplicates and standard deviations represented with error bars. For cytosine modification colour code, see **Figure 1**.

**Figure 6.**
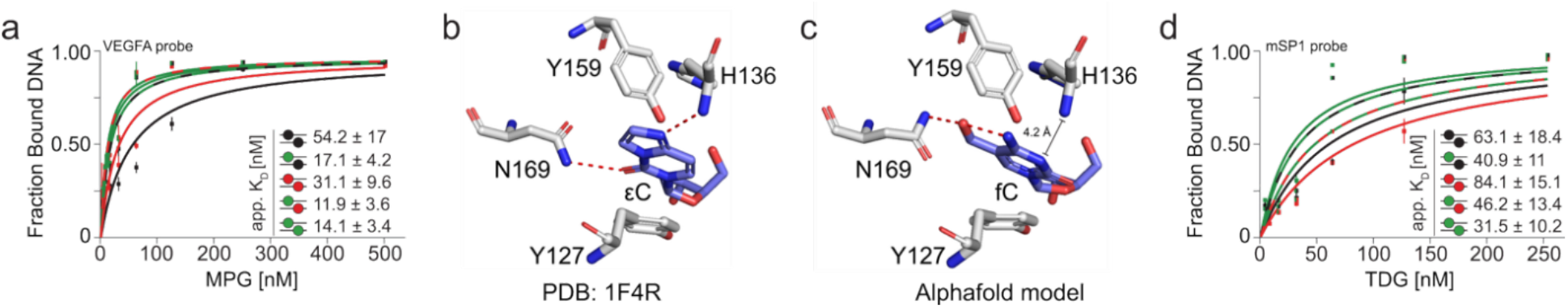
Validation of DNA repair proteins reading fC-modified CpG dyads with distinct symmetry preferences. (a) EMSA titration experiment with MPG binding to VEGFA probes. (b) Crystal structure of MPG (pdb: 1F4R) bound to DNA containing an εC lesion (blue). (c) AlphaFold model of MPG bound to DNA containing an fC nucleobase (blue). (d) EMSA titration experiments for TDG binding to mSp1 probes. Each experiment was done in duplicates and standard deviations represented with error bars. For cytosine modification colour code, see **Figure 1**.

We first studied the LZ-bHLH TF MAX that regulates cell growth, proliferation, and differentiation by forming dimers with itself and other LZ-bHLH TFs, most importantly MYC (49). MAX_2_ and MYC/MAX dimers bind the E-box (CACGTG) and E-box-like sequences, and are generally considered to be C/C CpG readers that are repelled by mC/mC CpGs (21,50). In our proteomics data, we found MAX to be enriched for fC/C over mC/mC, and it appeared as common fC/C reader throughout all human experiments (**Figure 3f**). Moreover, we also found MYC to be enriched for fC/C over fC/fC and fC/mC probes. Finally, other MAX dimerization partners of the LZ-bHLH TF class were enriched for fC-modified probes across different cell types and probe designs as well, such as MXD4, MXI1, MGA (**Figure 2k,o,q**), and the mouse proteins Mxi1 and Mga (**Figure 4o,r**). Indeed, MAX showed a preference for C/C and fC/C over other probe modifications in EMSA with the VEGFA probe (**Figure 5a**), in overall good agreement with the probe preferences in proteomics experiments (**Figure SI 26**). EMSA titration experiments using C/C and fC/C probes afforded very similar apparent (app.) K_D_s and revealed high affinity binding to both probes (**Figure SI 27a**). Given the presence of several E-box like sequences in the VEGFA probe (21), we also conducted EMSA titration experiments with the mSP1 probe containing only a single E-box-like sequence (CACGCC), and being modified only in CpG contexts. In agreement with the known C-selectivity of MAX_2_, we observed high affinity (app. K_D_=35 nm) for the probe containing C/C CpG dyads and low affinity for mC/mC (**Figure 5b**). Strikingly, MAX_2_ bound fC/C and fC/fC dyads with virtually the same affinity as C/C, whereas fC/mC was bound as weakly as mC/mC. This shows that high affinity E-box-like sequences exist in that MAX_2_ reads fC as strong as its cognate C base, making fC an activating mark compared to mC. It has previously been shown that MAX_2_ reads caC and antireads fC in the E-box sequence itself (50), whereas MAX_2_ and MYC/MAX can read hmC in other sequence contexts (21). Crystal structures show recognition of CpG and caCpG by MAX via a conserved arginine residue (R36 in MAX) (50,51), that may also be involved in fC recognition (**Figure 5c**). This combined data argues for a model in that fC and other oxi-mCs are read or antiread by MAX_2_ depending on the sequence context to possibly control its sequence-specificity and–given their conserved DNA binding motif–possibly other LZ-bHLH TFs. Interestingly, the catalytic domain of mammalian TETs shows pronounced selectivity for the E-box and certain E-box-like sequences, as well (52). In combination with the dysregulation/mutation of both MYC/MAX (53-55) and TETs (56) in cancer, the apparently complex interplay between these TFs with oxi-mC-containing target sequences warrants further investigation.

Next, we found another bHLH TF–the repressor HEY1–to be enriched for fC/C over mC/mC and fC/fC (**Figure 2o,q**). HEY1 is part of the Notch signaling pathway and regulates cell fate decisions by inhibiting differentiation and modulating proliferation during development and disease (57). EMSA with HEY1 and the mSP1 probe indeed revealed the fC/C dyad as a high affinity target, together with fC/fC and C/C dyads, whereas mC/mC and fC/mC dyads were bound with lower affinities (**Figure 5d**), suggesting that HEY1 is a C/C reader that is repelled by mC/mC, but that fC can confer high affinity unless it is paired with mC. This selectivity profile is similar to the one observed for MAX_2_ in the same probe context. Though there is no structure available for human HEY1 bound to DNA, the RR motif of MAX and other LZ-bHLH TFs containing the critical arginine involved in C recognition (R36 in MAX) is conserved in HEY1, which may account for the observed profile.

We also observed a series of RFX proteins to have interesting probe preferences in our proteomics data (these proteins share the conserved DNA-binding winged-helix (WH) domain (58)). In particular, we found RFX5 to be enriched for fC/mC and fC/fC over C/C (but not for fC/C over C/C), and it appeared as a common reader of mC/mC and fC/mC probes in all human experiments (**Figure 3f**). RFX5 is part of the RFX complex that binds to MHC Class II gene promoters and activates immune-related genes, and RFX5 mutations are disease-causing for the MHC class II deficiency disorder (59). RFX5 is an mC reader that binds to hmC/mC dyads with low and to hmC/hmC CpG dyads with even lower affinity, indicating that it is affected by modifications in both CpG strands (16,21). In EMSA with the VEGFA probe, the RFX5 DNA binding domain (DBD) showed a high affinity towards mC/mC followed by similar affinities for fC/mC and fC/fC and low affinities for C/C and fC/C probes (**Figure 5e** and **Figure SI 27b**), again in good agreement with the proteomics experiments (**Figure SI 26**). This again suggests that RFX5 is affected by modifications in both DNA strands. Moreover, unlike for hmC– where it acts as a general antireader (21) – it antireads fC in the fC/mC probe context, but its affinity is not further reduced for the fC/fC probe. This again fits to the overall picture that many mC readers tolerate fC better than hmC.

SIX2 is a homeobox TF that is involved in maintaining progenitor cells to prevent premature differentiation, most prominently during embryonic kidney development (60). We found SIX2 (along with its homolog SIX1) to be enriched for fC/fC over C/C, mC/mC and fC/C in proteomics experiments (**Figure SI 10i-j,q**). EMSA titration experiments with the mSP1 probe revealed that this protein acts as a selective reader of symmetric fC/fC dyads, discriminating against fC/C and C/C, and even more strongly against fC/mC and mC/mC dyads (**Figure 5f** shows app. K_D_s). The similar affinities for fC/C and C/C as well as fC/mC and mC/mC in the context of the asymmetric sequence contexts of CpGs in our probe prompted us to study the strand specificity of SIX2 by swapping fC to the respective antisense strand (**Figure 5g**). Indeed, EMSA revealed that SIX2 asymmetrically reads CpG dyads with strand-specific fC read-out in the case of fC/C and C/fC (compare **Figure 5f** and **h**). We also studied the SIX2 homolog SIX1 that showed very similar enrichment behavior in proteomics (**Figure SI 10i-j,q**). SIX1 plays a key role in embryonic development, particularly in organogenesis (61). The main difference between SIX1 and SIX2 lies in their tissue-specific expression and functional roles: SIX1 is more broadly expressed during embryogenesis and displays wide DNA sequence preferences, whereas SIX2 shows a more tissue-specific expression during embryogenesis (62). EMSA titration experiments with the mSP1 probe showed a related selectivity profile as observed for SIX2, yet with more refined dyad preferences (**Figure 5i**). SIX1 also read fC/fC dyads with highest affinity, followed by C-modified and mC modified dyads. However, in contrast to SIX2, this reader showed a slightly higher affinity for the latter dyads when a single fC was present. Again, a certain strand-specificity was observed when swapping fC between strands in the asymmetric dyads (compare **Figure 5i** and **j**).

We next analyzed FOXJ3 as representative of FOX proteins featuring a conserved DNA binding domain. FOXJ3 and many other FOX proteins showed specific probe preferences in all human proteomics experiments (**Figure 2** and **Figure SI 10,12**). FOXJ3 functions as a versatile, context-dependent TF that acts as tumor suppressor or promoter in neuroblastoma and breast cancer (63), respectively, and as transcriptional activator essential for meiotic gene expression in spermatogenesis (63). EMSA titration experiments revealed FOXJ3 as an fC reader with highest affinity for fC/fC, followed by fC/mC and fC/C, whereas C/C and particularly mC/mC were bound with weaker affinities (**Figure 5k**). FOXJ3 turned out to read fC in a strand-specific manner with reduced affinities for fC/C and fC/mC dyads when swapped (compare **Figure 5k** and **l**).

The tumor suppressor TP53 (p53) (64) has been identified as fC reader in previous mESC proteomics enrichments employing a short synthetic DNA probe containing fC/fC CpG dyads in a repetitive (CGA)_n_ context (16) that did not contain the p53 consensus sequence (which is composed of two consecutive half sites with the sequence: PuPuPu**C**(A/T)(T/A)**G**PyPyPy(65); particularly important residues in bold). Interestingly, we observed an enrichment of p53 for fC/C over C/C in mESC experiments (**Figure 4p**) and for fC/mC over fC/C and C/C probes in HEK293T experiments (**Figure 2k,m**). We conducted EMSA titration experiments with full-length p53 and the mSP1 probe containing two consecutive half sites with high similarity to the p53 consensus target sequence (C<u>AA</u>**<u>C</u>**<u>TT</u>**<u>G</u>**<u>CTC</u>/TT<u>A</u>**<u>C</u>**<u>A</u>C**<u>G</u>**<u>CCT</u>; underlined bases matching the consensus sequence), and containing a single CpG dyad with defined modifications (**Table S1**). EMSA titration experiments with this probe revealed slight preferences towards fC dyads, with affinities in the order of fC/C=fC/mC > fC/fC=C/C > mC/mC (**Figure SI 27c**).

We next studied BER proteins. MPG (N-methylpurine DNA glycosylase, also known as AAG) initiates BER by excising alkylated and deaminated purine bases from DNA (66). However, MPG also binds pyrimidine lesions such as 3,N_4_-ethenocytosine (εC) and 3-methylcytosine without being able to excise them (67). The lesions thus act as inhibitors that sequester MPG from actual target damage sites (68). Dependent on the sequence context and probe design, MPG has previously been identified as hmC (16) or fC reader (17), in both cases for symmetrically modified probes. We observed MPG as a common reader of any fC probe in human proteomics experiments (**Figure 3f-g**). In agreement with these observations (**Figure SI 26**), EMSA with the VEGFA probe revealed highest affinity towards fC/fC, but also the fC/mC and fC/C probes, followed by mC/mC and finally C/C (**Figure 6a**). MPG is a base flipping enzyme that recognizes an inhibiting εC base with similar affinity as important substrate lesions such as εA. In crystal structures, εC (and εA) are recognized by stacking interactions with Y127 on one side and Y159 and H136 on the other side, whereas N169 donates an h-bond to the O2 of εC (**Figure 6b**). The main chain amide of H136 donates an h-bond to the N6 of εA or N4 of εC (3.2 Å distance for the latter), in both cases being an h-bond acceptor rather than a donating amino group as in the respective canonical base, enabling selective recognition of the damaged base (69). An alphaFold model of MPG bound to an fC-containing DNA shows a highly similar overall recognition of fC and εC (**Figure 6c**). However, the distance between the H136 amide and the fC N3 atom is 4.2 Å, and thus beyond the expected distance for a hydrogen bond. In contrast, the side chain of N169 is rotated to form an h-bond to the 5-formyl group of fC, which may explain the observed higher affinity for fC over undamaged C (**Figure 6c**). We also conducted EMSA using the mSP1 probe and observed a different affinity profile, with mC/mC being bound with highest affinity, and fC being bound with different affinities dependent on the C state in the opposite dyad strand (**Figure SI 27d**). Moreover, when fC was put into yet another sequence context within the same mSP1 probe by swapping the dyad strand, we again observed an altered affinity in the case of C/fC versus fC/C dyads (**Figure SI 27e**). In the light of known sequence dependencies of MPG (70,71) and previous conflicting oxi-mC reading observations for different probe sequences (16,17), this data hints at a complex interdependence of sequence and fC recognition by MPG that requires further investigation to understand fC’
ss potential roles as an inhibitor of MPG repair functions.

Next, we studied TDG, the BER enzyme excising thymine from G:T mismatches as well as fC and caC from CpG dyads to initiate the BER pathway (**Figure 1a**) (7,72). TDG has been found to preferentially bind fC in several proteomics and other studies and it appeared as common reader of all fC-containing probes in our human and mouse proteomics experiments (**Figure 3f** and **5b,d,e**). In agreement with previous EMSA experiments employing probes with symmetrically modified CpG dyads (16), we observed the highest complex formation for fC/fC dyads, followed by C/C and mC/mC dyads. So far unknown, we found that the asymmetric fC/C and fC/mC dyads were bound with lower affinity than fC/fC dyads. (**Figure 6d**, see **Figure SI 27f** for strand swapping experiment). Next, the apurinic/apyrimidinic endonuclease 1 (APEX1) initiates BER as the enzyme cleaving apurinic/apyrimidinic sites in damaged DNA (e.g., generated by TDG), but also regulates TFs via redox modulation and participates in RNA processing (73). As TDG, it was enriched for fC-modified probes in human and mouse experiments (**Figure 3f** and **5b,d,e**). EMSA titration experiments with the mSP1 probe revealed only subtle affinity differences, which may be explained by an indirect recruitment of APEX1, e.g., via its interaction partner TDG (**Figure SI 27g**). We obtained similar results for XRCC1 (X-ray repair cross-complementing protein 1), another component of the BER pathway that interacts with TDG (**Figure SI 27h**) (37).

## DISCUSSION

We have employed an MS-based approach to generate the first proteome-wide interaction profiles of symmetric and asymmetric fC modifications in DNA. We systematically and comparatively studied human and mouse nuclear proteomes employing three different DNA enrichment probes derived from human and mouse promoter regions containing CpG dyads with five symmetric and asymmetric C, mC and fC-modification combinations. For all cell lines, the number of proteins reading fC-modified probes exceeds that of proteins reading the canonical C/C and mC/mC probes combined. This suggests, from a biochemical perspective, that fC modifications are important protein interaction sites in the genome, with implications for their role in chromatin regulation. fC/fC probes thereby attracted a high number of proteins, which is in agreement with previous studies (16). However, we discover that a similar or even higher number of proteins are attracted by the two asymmetric fC probes. In the light of previous symmetry-resolved proteomics data for hmC (21), our new data support the overall view that fC is a more permissive and often even preferred modification for many proteins as compared to hmC, which is considered to be a more repelling mark (16-18,74). The relative reader numbers for the three individual fC probe designs overall vary from cell line to cell line, and we observe a significant sequence- and tissue-specificity of readers. This suggests that many more fC readers could be identified by profiling additional tissues and using additional probe designs. However, the overlap between readers of all five probes seems less dependent on tissue and probe sequence. All three fC modifications attract a high number of unique readers and overlap only partially among each other and with C/C and mC/mC probes. We could extract a set of common readers for each modification that were enriched for all human tissues and employed probe sequences. For C/C and mC/mC probes, this included the canonical C and mC readers identified in previous studies. In contrast, all fC probes attracted BER proteins along with TFs and other chromatin regulators, often being specific for each of the three fC probes. We selected eleven reader candidates for detailed evaluation of their modification preferences in vitro, and observe symmetry-specific (and partially also strand-specific) fC reading preferences in most cases. These proteins include TFs with central roles in development and disease, such as MAX_2_, HEY1, RFX5, SIX1, SIX2, TP53 and FOXJ3. For several of these readers, related family members with high sequence similarity or even structurally conserved DNA recognition domains display similar probe preferences in proteomics experiments, suggesting broader implications. For example, we found that many MAX dimerization partners and other bHLH TFs were enriched for fC-modified probes, including MXD4, MXI1, MGA and the mouse proteins Mxi1 and Mga. Moreover, we observe that MAX reads or antireads fC (and hmC (21)) in a sequence-dependent manner, suggesting that oxi-mC may control its sequence-specificity (an observation made previously for several other TFs (22)). The preference of the TET catalytic domain for the E-box and E-box-like sequences (52) and the mutation of both TETs and bHLH-LZ TFs in many cancers (53,56) hint at a possible, complex involvement of oxi-mC in cancer development that deserves further investigation. We also validate BER proteins and find TDG and MPG to exhibit fC symmetry preferences, in the latter case again with either fC reading or anti-reading dependent on the sequence context.

For evaluating the actual physiological relevance of the identified interactions including their tissue-specificity, it will be required to correlate sequencing-based high resolution genomic maps for specific readers with high resolution maps for individual CpG dyad modifications. For example, during early embryonic development, fC shows specific enrichment in the paternal pronuclei (46) and acts as activating mark in *Xenopus* oocytes (15). fC also seems to play a role in nucleosome positioning (13) and impairs RNA polymerase II elongation and fidelity (14). It is however not understood, in which CpG dyad contexts fC occurs in these situations. Unfortunately, whereas CpG dyad-resolved hmC sequencing methods have been reported very recently (75-78), comparable methods are not yet available for fC (20). Altered oxi-mC levels are also a hallmark of many cancers, with TET2 being one of the most frequently mutated genes in hematopoietic malignancies (56). In addition, IDH1/2 mutations and other causes of altered concentrations of citric acid cycle intermediates (2-oxoglutarate, 2-hydroxyglutarate, fumarate) modulate TET activity in such cancers. fC has further been shown to occur at altered levels in breast and prostate cancer as well as in myeloid malignancies, with implications for use as biomarker (79-81). Our reader catalogue will thus help studying the functional connection between fC and malignant processes with resolution for different fC dyad symmetries.

In conclusion, our study provides a foundation for dissecting molecular mechanisms of chromatin regulation that may be governed by symmetric and asymmetric fC modifications, with particular importance for development and cancer.

## Supporting information

Table S6. JASPAR motif analysis

Table S7. Significantly enriched proteins of pulldown with HEK293T and VEGFA probe

Table S8. Significantly enriched proteins of pulldown with HEK293T and hSP1

Table S9. Significantly enriched proteins of pulldown with HeLa and VEGFA probe

Table S10. Common readers_Gene Ontology Terms

Table S11. Significantly enriched proteins of pulldown with mESC and mSP1probe

Table S12. mESC readers_Gene Ontology Terms

## SUPPLEMENTARY DATA

Supplementary Data are available online: **Table SI 1-12 and Figure SI 1-27**.

## ACKNOWLEDGEMENT

We thank the Faculty of Chemistry and Chemical Biology of the TU Dortmund University, the Max-Planck Society and the International Max-Planck Research School for Living Matter for continuous support. We thank the members of the group for support. We thank Dr Christian Schöter for mouse embryonic stem cells (E14TG2a) and LIF. Funded by the Deutsche Forschungsgemeinschaft (SU726/9-1). Funded by the CANTAR program “Netzwerke 2021”, an initiative of the Ministry of Culture and Science of the State of Northrhine Westphalia.

## AUTHOR CONTRIBUTIONS

Z.V.C. developed methods, conducted wet lab experiments, analyzed data and conducted modelling experiments. L.E. developed methods. N.S. cloned plasmids and purified proteins. S.E. purified proteins. T.B. supervised MS experiments and analyzed MS data. D.S and Z.V.C. wrote the manuscript with the help of all other authors. D.S. analyzed data and conceived and supervised the study.

