## Supporting Information for "Proteomics-Based Discovery of Symmetry-Specific Readers and Antireaders of 5-Formylcytosine in Mammalian DNA"

This PDF file includes:

Supplementary Table **1 to 5**

Supplementary Fig. **1 to 27**

**Table S1.** DNA probes

| Description | Sequence 5′>3′ |
|---|---|
| VEGFA probe | TTTCCAAAGCCCATTCCCTCTTTAGCCAGAGCCGGGGTGTGCAGACGGCAGTCACTAGGGGGCG CTCGGCCACCACAGGGAAGCTGGGTGAATGGAGCGAGCAGCGTCTTCGAGAGTGAGGACGTGTG TGTCTGTGTGGGTGAGTGAGTGTGTGCGTGTGGGGTTGAGGGCGTTGGAGCGGGGAGAAGGCCA GGGGTCACT |
| hSP1 probe | GGGGGTAAGATTTGAGAGGTACTTTATAGGGGCAGTTAAATGAAGACGCAAACAAGTCCTAGTG TTGATGCGGAACTGCGCGCCGAATGCCTTGGCTCTGACACCTGTTGAGCTGCAGGACTCCGCTA AAGCGTCCCACCTAATGACTGTAACAACGTCCCCTGAGGAGGGCCAATATGGCGACGGTCTCCT CTTGGCATAGCCCTCTTCCCTCCC |
| mSP1 probe | AAGTACGCAACTTGCTCTTACACGCCTCAGCGAGAGAGCGAGTCCTACCA |

**Table S2.** Oligonucleotides used for probe generation.

| Oligo number | Description | Sequence 5′>3′ |
|---|---|---|
| o4681 | VEGFA frw | [BtnTg]TTTCCAAAGCCCATTCCCT |
| o4645 | VEGFA frw with FAM | [BtnTg]TTTCCAAAGCCCA[FAM]TCCCT |
| o4666 | VEGFA rev | AGTGACCCCTGGCCT |
| o6120 | hSP1 frw | [BtnTg]GAGAGGTACTTTATAGGGGCAG |
| o6119 | hSP1 rev | GGGAGGGAAGAGGGCTATG |
| o6439 | mSP1 frw, C | [FAM]AAGTACGCAACTTGCTCTTACACGCCTCAGCGAGAGAGCGAGTCCTACCA [BtnTg] |
| o6440 | mSP1 rev, C | TGGTAGGACTCGCTCTCTCGCTGAGGCGTGTAAGAGCAAGTTGCGTACTT |
| o6441 | mSP1 frw, mC | [FAM]AAGTAXGCAACTTGCTCTTACAXGCCTCAGXGAGAGAGXGAGTCCTACCA [BtnTg], X=mC |
| o6442 | mSP1 rev, mC | TGGTAGGACTXGCTCTCTXGCTGAGGXGTGTAAGAGCAAGTTGXGTACTT, X=mC |
| o6443 | mSP1 frw, fC | [FAM]AAGTAXGCAACTTGCTCTTACAXGCCTCAGXGAGAGAGXGAGTCCTACCA [BtnTg], X=fC |
| o6444 | mSP1 rev, fC | TGGTAGGACTXGCTCTCTXGCTGAGGXGTGTAAGAGCAAGTTGXGTACTT, X=fC |

**Table S3.** Oligonucleotides used for CDS amplification of full-length proteins and Gibson assembly. The overhangs are indicated in lowercase letters.

| Name | Description | Sequence |
|---|---|---|
| o6145_ZeC | HEY1 frw | gaaaatctttattttcagtctctcAAGCGAGCTCACCCCGAG |
| o6146_ZeC | HEY1 rev | cagtggtggtggtggtggtgctcAAAAGCTCCGATCTCCGTCC |
| o6207_ZeC | SIX2 frw | gaaaatctttattttcagtctctcTCCATGCTGCCCACCTTC |
| o6208_ZeC | SIX2 rev | cagtggtggtggtggtggtgctcGGAGCCCAGGTCCACGAG |
| o6385_ZeC | SIX1 frw | gaaaatctttattttcagtctctcTCGATGCTGCCGTCGTTTG |
| o6386_ZeC | SIX1 rev | cagtggtggtggtggtggtgctcacttccaccGGACCCCAAGTCCACCAGAC |
| o6363_ZeC | FOXJ3 frw | gaaaatctttattttcagtctctcGGTTTGTATGGACAGGC |
| o6364_ZeC | FOXJ3 rev | cagtggtggtggtggtggtgctcacttccaccCACAATTGAATCCCAATC |
| o6299_ZeC | TP53 frw | gaaaatctttattttcagtctctcGAGGAGCCGCAGTCAGATCC |
| o6300_ZeC | TP53 rev | cagtggtggtggtggtggtgctcGTCTGAGTCAGGCCCTTCTG |
| o5748_ZeC | MPG frw | gaaaatctttattttcagtctctcATGGTCACCCCCGCTTTG |
| o5749_ZeC | MPG rev | cagtggtggtggtggtggtgctcGGCCTGTGTGTCCTGCTCAG |
| o5774_ZeC | TDG frw | gaaaatctttattttcagtctctcATGGAAGCGGAGAACGCG |
| o5775_ZeC | TDG rev | cagtggtggtggtggtggtgctcAGCATGGCTTTCTTCTTCCTG |
| o6281_ZeC | APEX1 frw | gaaaatctttattttcagtctctcCCGAAGCGTGGGAAAAAG |
| o6282_ZeC | APEX1 rev | cagtggtggtggtggtggtgctcCAGTGCTAGGTATAGGGTG |
| o5832_ZeC | XRCC1 frw | gaaaatctttattttcagtctctcATGCCGGAGATCCGCCTC |
| o5833_ZeC | XRCC1 rev | cagtggtggtggtggtggtgctcGGCTTGCGGCACCACCCC |

**Table S4.** Common readers of VEGFA and hSP1 probes

| C/C | Count | Proteins |
|---|---|---|
| Common readers | 36 | INO80E ASH2L KMT2B TFPT MEN1 NFRKB RMI1 SMAD2 USF2 NFIC BLM MYC BCOR MAX THAP1 NFIA ACTR8 E2F3 TBRG1 NAIF1 PCGF1 RMI2 TERF2IP ELF2 NFIX KAT6B L3MBTL3 KMT2A HEY1 XBP1 TFAP2A KDM2B USF1 TFAP4 RBPJ FLYWCH1 |
| VEGFA probe specific | 71 | CLOCK ARNT CETN3 KDM5D NSMCE3 ATF6B ATF6 PROX1 NELFE CDCA7 ORC3 MXD4 DGCR8 HMBOX1 BMAL1 PPP1R10 NUP155 KPNA4 NKX2-5 NKX2-2 SETD2 SMAD4 CENPQ SMARCC2 POLR2J ZIC3 CBFB CEBPB KMT2D YY1AP1 DRAP1 ESRRA RBBP5 CDK9 TFE3 FAAP24 ZNF609 MED10 CENPB NOLC1 HERC2 CCNL1 CABIN1 RSBN1 RUNX3 NOL8 MIS18BP1 H2BC26 NUFIP1 HES1 BRD8 RUNX1 ARID1A SUV39H1 MED17 ELP1 PPWD1 TOX4 NKX2-4 PBX1 MXI1 AHR SNRNP35 CDK13 INTS2 TAF1B NRF1 BUD23 NET1 CDYL RNF169 |
| hSP1 probe specific | 74 | PKNOX1 LCOR POLR2B PSMA4 REXO4 DDX51 MTF2 NFYC CFL1 TIGD7 LGALS7B HOXD9 KDM1A BMS1 SFN HNRNPH2 MAFG TFAP2C ZMYND11 PPHLN1 EXOSC4 NIP7 APEX1 INO80 KAT6A ILF3 TRIP12 EXOSC1 SNRPC CPSF7 KDM2A SCAF11 TCF12 SNU13 TERF2 MPHOSPH10 SUZ12 NOL10 RPP25L MYPOP CASP14 EED WDR18 E2F6 NUDT21 CENPX CBX1 ATF3 RBM45 NFIB SF3A1 ADAR LAS1L UCHL5 FOSB ELF4 RPF2 TFDP1 MTA1 ZNF22 FBXW11 RING1 EXOSC7 RRS1 PPAN ZNF280C APOBEC3C EZH2 EXOSC3 RNF2 PARP1 ING5 HAND1 NACA |

| fC/C | Count | Proteins |
|---|---|---|
| Common readers | 25 | CDCA7L FOXD2 NFRKB RMI1 MAX PTBP2 ACTR8 ATF3 RBM45 L3MBTL3 TFDP1 KDM2B NTHL1 TDG PCGF6 APEX1 THAP1 MPG TCF12 NAIF1 RMI2 HEY1 TFAP2A APOBEC3C TIGD1 |
| VEGFA probe specific | 105 | ATF6B ATF6 INO80E BMAL1 ZFHX4 NKX2-5 PNKP SUB1 PURB RFX5 SMAD2 CBFB ZMYM2 KMT2D BLM DRAP1 RRP8 RFC2 PAX1 BRCA2 SIX2 L3MBTL2 SSBP3 UBN1 CENPB NOLC1 RUNX3 MECP2 TBRG1 TERF2 CENPS MBD4 TERF2IP E2F6 HES1 ISL1 SMAD3 PAX3 PAX6 DLX2 UCHL5 VRK3 LDB1 FOSB PCBP2 TOX4 MXI1 ASF1A RBPJ PURA RFXAP CREB1 FANCM ARNT PARP2 RPA2 JUNB MXD4 APTX VAX2 PPP1R10 SKI RPA1 NKX2-2 SMAD4 XRCC6 XRCC1 PAX9 KLF5 TBX2 PAX2 MYC ESRRA LIG3 RFC1 RFC4 FAAP24 TBX3 CABIN1 RFC5 HIRA ISL2 SMARCAL1 DCLRE1B CENPX KLHL7 RUNX1 NR2C2 POLB BARX1 DR1 CREM NR2F2 XBP1 RYBP NKX2-4 PBX1 EMX1 AHR TEAD4 TDP1 RPA3 NR2F6 NKX1-2 PARP1 |
| hSP1 probe specific | 134 | SMNDC1 DDX51 DDX1 ATRX RAE1 MTF2 FYTTD1 H3C12 ZNF146 ADARB1 MAFG PHIP PPHLN1 MRTO4 EXOSC4 NIP7 CD2BP2 MBD3 SCAF1 EXOSC1 NFIA MCRS1 SNRPC NPM3 NMNAT1 FOXC2 EBNA1BP2 H3C13 RAI1 MPHOSPH10 ATAD2 ZMAT2 RPP25L DDX31 EED WDR18 NFIX CBX1 TAF5 ZCRB1 SUV39H1 RSL24D1 POLR2C NFIB DDX27 TRIM27 DNMT3A CHRAC1 NUP205 RING1 TFAP4 TBX1 EXOSC7 ZNF644 RRS1 FLYWCH1 ZNF280C CHD2 EXOSC3 FOXD1 ING5 SRPK2 VIRMA LCOR POLR2B FOXC1 REXO4 CDCA7 TAF15 KAT8 SRRT BMS1 BAZ2A SPATS2L HNRNPH2 MPHOSPH6 RSF1 KHDRBS1 HMG20A TFAP2C RTRAF NPM1 NXF1 NCBP2 CEBPB NFIC KAT6A BCOR RANBP2 GTF3C4 ILF3 H2AX TRIP12 KDM2A SCAF11 MACROH2A1 SCAF4 CLASRP EXOSC5 RCC1 TBX18 RBM5 SNU13 EXOSC8 PCGF1 H2BC26 NOL10 RRP9 ELF2 NGDN SMARCE1 SATB1 KAT6B MACROH2A2 BRD1 CDCA5 ADAR LAS1L POP1 RPF2 MTA1 CEBPA ZNF22 FBXW11 SNRNP35 USF1 NSA2 SRSF12 SURF6 FOXG1 RTCB NEDD8 PPAN CDCA8 |

| mC/mC | Count | Proteins |
|---|---|---|
| Common readers | 28 | FOXC1 FOXF2 ZFHX4 ATF2 RFX5 RECQL FOXB1 SSBP3 FOXC2 ATF7 MECP2 MBD2 RMI2 ISL2 MBD4 ISL1 MBD1 RBM45 DLX2 RBM14 ZHX1 FOXK1 LDB1 MSH2 TFAP2A CREB5 FBXO11 FOXA1 |

| | | |
|---|---|---|
| VEGFA probe specific | 72 | PRPF40A VIRMA ACIN1 DACH2 PARP2 NSMCE3 ZNF207 NELFE RPA2 DGCR8 JUNB CLK4 PPP1R10 UHRF2 CLK2 TBL1X NKX2-5 SETD2 RPA1 CSTF1 PPHLN1 TBL1XR1 SUB1 RMI1 SMARCC2 RBM25 POLR2J RFX7 TOE1 YY1AP1 BLM ESRRA ILF3 HOXA13 RFX1 FAAP24 SRSF7 ZNF609 XRCC4 CABIN1 RSBN1 SRSF3 HOXB13 H3C13 TET1 CENPS ZNF668 SMARCAL1 FOXK2 RFXANK WWP1 API5 HOXD13 SUV39H1 SPEN BAP1 CLK3 PPWD1 WTAP DNMT3A NKX2-4 SF1 CDK13 RBPJ RFX2 SRSF10 RPA3 NET1 RFXAP RBBP6 FANCM TFCP2 |
| hSP1 probe specific | 48 | FOXD2 VAX2 MEIS2 HOXB5 SATB2 NUP153 FOXO3 HOXB8 FOXP2 UHRF1 DLX1 MEIS1 MIER1 TBX2 FOXJ3 TAF10 GATAD2A ZNF687 TEAD1 DLX5 ALX4 TBX3 ARID5B GBX2 PBX2 FOXP1 FOXO1 CHD4 HOXA4 ZFHX3 PBX3 SATB1 SSBP2 BARX1 LMX1B HOXB4 MNX1 EN2 SSBP4 PBX1 ALX1 TBX1 TEAD4 FOXG1 PARP1 FOXD1 DLX6 PDX1 |

| **fC/mC** | **Count** | **Proteins** |
|---|---|---|
| Common readers | 28 | MPG FOXC2 NTHL1 FOXC1 TDG TBX18 MECP2 MBD2 RMI2 ISL2 FOXO1 FOXF2 MBD4 FOXK2 ISL1 MBD1 RFX5 RMI1 BAP1 ZHX1 FOXK1 LDB1 BLM FOXB1 FOXG1 FOXA1 RFXAP SSBP3 |
| VEGFA probe specific | 42 | DACH2 CABIN1 NSMCE3 ORC3 HOXB13 DGCR8 JUNB NAIF1 VAX2 NUP155 RFXANK TP53 API5 ZFHX4 DACH1 KLHL7 PAX3 POLB PURB SUB1 TOE1 APEX1 TBX2 SIX1 FOXJ3 PBX1 EMX1 YY1AP1 DRAP1 ASF1A SAFB2 CDK13 SAFB RBPJ RFX2 SRSF10 SIX2 RFX1 HOXA13 FAAP24 NET1 TBX3 |
| hSP1 probe specific | 39 | CDCA7 DDX1 ASXL2 FOXD2 FOXP1 H2BC26 EYA1 FOXO4 SATB2 ZFHX3 FOXO3 RTRAF RBM45 PAX6 SATB1 UHRF1 SSBP2 TBL1XR1 LMX1B PAX9 MSH2 NKX3-2 SSBP4 MTA1 CEBPA TFAP2A TBX1 TEAD4 ERF MAX NCOR1 RTCB TRIP12 TIGD1 TEAD1 FOXD1 FBXO11 KDM5A ING5 |

| **fC/fC** | **Count** | **Proteins** |
|---|---|---|
| Common readers | 34 | NTHL1 FOXC1 TDG RBM3 VAX2 FOXF2 PAX9 APEX1 CEBPB SIX4 FOXB1 PAX1 THAP1 SIX2 MPG FOXC2 TBX18 MECP2 FOXO1 MBD4 ETV3 FOXK2 TAF5 KLHL7 PAX3 POLB BAP1 FOXK1 SIX1 CEBPA TBX1 ERF FOXG1 TIGD1 |
| VEGFA probe specific | 70 | CETN3 NELFE TAF3 ORC3 PSPC1 DGCR8 JUNB KIN STRAP RPF1 NUP155 TAF8 TAF12 GTF2A1 DAZAP1 CENPH TAF4 SETD2 PCGF6 CENPQ TAF9B RFX5 HNRNPD SMARCC2 TAF6 POLR2J EOMES TBX2 DRAP1 TAF10 PTBP2 L3MBTL2 TBX3 NONO MED10 TAF7 TAF11 HERC2 RSBN1 TAF9 H3C13 CIRBP PRMT5 ZNF668 H2BC26 KNL1 RFXANK E2F6 MED14 GTF2A2 BRD8 ATF3 SUV39H1 REPIN1 MED17 RBM14 SSX2IP TRIM27 ELP1 H1-6 PPWD1 TFDP1 RYBP MGA TAF1 CDK13 INTS2 TAF1B APOBEC3C RFXAP |
| hSP1 probe specific | 57 | PARP2 DDX1 TIGD7 FAM98B APTX FOXD2 EYA1 CHAF1A HOXB5 NUP153 FOXO3 RTRAF PNKP TBL1XR1 DLX1 PAX2 FOXJ3 LHX2 LIG3 RFC1 RFC2 RFC4 TTF1 DLX5 SNRPC HHEX ALX4 SSBP3 FOXF1 RFC3 PCNA RBBP4 CHAF1B GBX2 RFC5 CDC6 FOXP1 ISL2 FOXO4 ISL1 VAX1 RSBN1L PAX6 SSBP2 LMX1B DDB2 MNX1 EN2 LDB1 SSBP4 ALX1 RTCB CHTF18 FOXD1 FBXO11 DLX6 PDX1 |

**Table S5.** Common readers of HEK293T and HeLa

| C/C | Count | Proteins |
|---|---|---|
| Common readers | 20 | ARNT ATF6 RMI1 CBFB CEBPB BLM BCOR ESRRA MAX FAAP24 RUNX3 NAIF1 RMI2 TERF2IP RUNX1 L3MBTL3 KMT2A TFAP2A KDM2B AHR |
| HEK293T specific | 87 | CLOCK CETN3 KDM5D NSMCE3 ATF6B PROX1 NELFE CDCA7 ORC3 MXD4 DGCR8 INO80E HMBOX1 BMAL1 ASH2L KMT2B PPP1R10 NUP155 TFPT KPNA4 NKX2-5 NKX2-2 SETD2 SMAD4 CENPQ MEN1 NFRKB SMARCC2 POLR2J SMAD2 ZIC3 USF2 KMT2D NFIC YY1AP1 MYC DRAP1 RBBP5 THAP1 CDK9 TFE3 NFIA ZNF609 MED10 CENPB ACTR8 NOLC1 HERC2 CCNL1 E2F3 CABIN1 RSBN1 TBRG1 PCGF1 NOL8 MIS18BP1 H2BC26 NUFIP1 ELF2 HES1 BRD8 NFIX ARID1A SUV39H1 MED17 KAT6B ELP1 PPWD1 HEY1 TOX4 XBP1 NKX2-4 PBX1 USF1 MXI1 SNRNP35 TFAP4 CDK13 INTS2 RBPJ FLYWCH1 TAF1B NRF1 BUD23 NET1 CDYL RNF169 |
| HeLa specific | 51 | LIG4 RBFOX1;RBFOX2 TOP2B RPS3A ACIN1 DPY30 CCAR2 SRP68 RPA2 SSRP1 CTR9 AHRR EIF6 KPNB1 ZMYND11 UHRF1 SUB1 RPLP0 PAXX SERBP1 RACK1 KDM2A HNRNPL MACROH2A1 ATF1 CETN2 TCF12 NCL H3C13 TERF2 CENPS SMARCAL1 RPS10 HOXD12 BANP CENPX H2AC19;H2AC20 XPC RBM45 TRA2B MSH2 RAD23B HNRNPK PCBP2 RPL5 RUNX2 RPA3 HNRNPUL2 MYO1C FANCM NACA |

| fC/C | Count | Proteins |
|---|---|---|
| Common readers | 22 | ARNT NTHL1 CDCA7L TDG RPA2 PURB RMI1 APEX1 CBFB BLM MAX MPG NAIF1 CENPS RMI2 MBD4 TERF2IP RUNX1 RBM45 AHR PURA FANCM |
| HEK293T specific | 108 | PARP2 ATF6B ATF6 MXD4 JUNB INO80E APTX FOXD2 VAX2 BMAL1 PPP1R10 ZFHX4 SKI NKX2-5 PNKP NKX2-2 RPA1 SMAD4 PCGF6 XRCC6 NFRKB XRCC1 SUB1 KLF5 PAX9 RFX5 SMAD2 PAX2 TBX2 ZMYM2 KMT2D MYC DRAP1 RRP8 ESRRA LIG3 RFC2 RFC1 THAP1 PAX1 BRCA2 RFC4 PTBP2 SIX2 L3MBTL2 FAAP24 TBX3 SSBP3 UBN1 ACTR8 CENPB NOLC1 TCF12 CABIN1 RFC5 RUNX3 MECP2 TBRG1 HIRA TERF2 ISL2 SMARCAL1 DCLRE1B E2F6 HES1 ISL1 CENPX SMAD3 KLHL7 ATF3 PAX3 PAX6 NR2C2 POLB DLX2 BARX1 L3MBTL3 UCHL5 DR1 CREM VRK3 LDB1 FOSB NR2F2 TFDP1 PCBP2 HEY1 TOX4 XBP1 RYBP NKX2-4 TFAP2A EMX1 KDM2B PBX1 MXI1 ASF1A TEAD4 TDP1 RBPJ RPA3 NR2F6 APOBEC3C TIGD1 NKX1-2 PARP1 RFXAP CREB1 |
| HeLa specific | 24 | RBFOX1;RBFOX2 SRP68 RBM3 AHRR ORC1 KPNB1 MAFG HNRNPD HNRNPA0 HNRNPAB SYNCRIP CEBPB IGF2BP2 HNRNPL ATF1 H3C13 CIRBP TFAP2D HNRNPK FOXE1 RPL5 RUNX2 HNRNPUL2 MYO1C |

| mC/mC | Count | Proteins |
|---|---|---|
| Common readers | 29 | FOXC2 ACIN1 ATF7 SRSF3 RPA2 MBD2 CENPS RMI2 MBD4 RFXANK UHRF2 ATF2 MBD1 SUB1 RFX5 RMI1 RECQL FOXK1 MSH2 BLM RBPJ SRSF10 RFX2 RFX1 FAAP24 FOXA1 RFXAP SRSF7 FANCM |
| HEK293T specific | 71 | PRPF40A VIRMA DACH2 FOXC1 CABIN1 PARP2 NSMCE3 ZNF207 RSBN1 NELFE MECP2 DGCR8 HOXB13 JUNB H3C13 TET1 CLK4 ZNF668 ISL2 FOXF2 SMARCAL1 PPP1R10 FOXK2 WWP1 API5 ISL1 ZFHX4 CLK2 TBL1X NKX2-5 HOXD13 SETD2 SUV39H1 RBM45 RPA1 CSTF1 PPHLN1 TBL1XR1 DLX2 RBM14 SPEN SMARCC2 RBM25 BAP1 POLR2J RFX7 ZHX1 TOE1 CLK3 LDB1 PPWD1 WTAP DNMT3A NKX2-4 TFAP2A YY1AP1 ESRRA SF1 CDK13 FOXB1 ILF3 CREB5 RPA3 HOXA13 FBXO11 NET1 ZNF609 XRCC4 RBBP6 SSBP3 TFCP2 |

| | | |
|---|---|---|
| HeLa specific | 24 | FOXF1 CETN2 SRSF2 SAP18 HIRA RNPS1 TLX3 CHD4 CENPX MTA2 ARL6IP4 UHRF1 C1QBP TIA1 RAD23B GATAD2B FOXE1 THRAP3 CHD3 GATAD2A PNN SRSF6 PUF60 UBN1 |

| fC/mC | Count | Proteins |
|---|---|---|
| Common readers | 14 | MPG FOXC2 TDG NAIF1 MBD2 MBD4 RFXANK MBD1 PURB RFX5 FOXK1 BLM FOXA1 RFXAP |
| HEK293T specific | 56 | NTHL1 DACH2 FOXC1 CABIN1 NSMCE3 TBX18 MECP2 ORC3 HOXB13 DGCR8 JUNB RMI2 ISL2 VAX2 FOXO1 FOXF2 NUP155 FOXK2 TP53 API5 ISL1 ZFHX4 DACH1 KLHL7 PAX3 POLB SUB1 RMI1 BAP1 ZHX1 TOE1 APEX1 LDB1 TBX2 SIX1 FOXJ3 PBX1 EMX1 YY1AP1 DRAP1 ASF1A SAFB2 CDK13 FOXB1 SAFB RBPJ RFX2 SRSF10 FOXG1 SIX2 RFX1 HOXA13 FAAP24 NET1 TBX3 SSBP3 |
| HeLa specific | 17 | FOXF1 CBX5 PARP2 TLX3 CHD4 ORC1 HOXC10 MTA2 UHRF2 UHRF1 TIA1 HNRNPAB TRA2B FOXE1 GATAD2A PURA PUF60 |

| fC/fC | Count | Proteins |
|---|---|---|
| Common readers | 17 | NTHL1 TDG RBM3 RFX5 HNRNPD APEX1 CEBPB L3MBTL2 TBX3 MPG FOXC2 CIRBP MBD4 E2F6 ATF3 FOXK1 RFXAP |
| HEK293T specific | 87 | CETN3 FOXC1 NELFE TAF3 ORC3 PSPC1 DGCR8 JUNB KIN STRAP VAX2 RPF1 FOXF2 NUP155 TAF8 TAF12 GTF2A1 DAZAP1 CENPH TAF4 SETD2 PCGF6 CENPQ TAF9B PAX9 SMARCC2 TAF6 POLR2J EOMES TBX2 SIX4 DRAP1 FOXB1 TAF10 PAX1 THAP1 PTBP2 SIX2 NONO MED10 TAF7 TAF11 HERC2 RSBN1 TBX18 MECP2 TAF9 H3C13 PRMT5 ZNF668 FOXO1 H2BC26 KNL1 ETV3 FOXK2 RFXANK MED14 GTF2A2 BRD8 TAF5 KLHL7 PAX3 SUV39H1 REPIN1 POLB MED17 RBM14 SSX2IP TRIM27 BAP1 ELP1 H1-6 PPWD1 SIX1 TFDP1 CEBPA RYBP MGA TBX1 TAF1 ERF CDK13 INTS2 FOXG1 TAF1B APOBEC3C TIGD1 |
| HeLa specific | 37 | RBFOX1;RBFOX2 HNRNPDL PARP2 RPA2 H1-10 HNRNPR WDR5 CHAF1A ORC1 MTA2 CEBPD JDP2 CSRP1 CBX3 HNRNPA0 HNRNPAB SYNCRIP FEN1 RFC2 RFC4 HNRNPU RFC3 FOXF1 ATF1 HNRNPA3 CHAF1B SFPQ RFC5 BAZ1B SUMO2 HNRNPA2B1 RCC2 HNRNPK FOXE1 EEF1D HNRNPUL2 NACA |

**As attached excel file:**

Table S6. JASPAR motif analysis

Table S7. Significantly enriched proteins of pulldown with HEK293T and VEGFA probe

Table S8. Significantly enriched proteins of pulldown with HEK293T and hSP1

Table S9. Significantly enriched proteins of pulldown with HeLa and VEGFA probe

Table S10. Common readers_Gene Ontology Terms

Table S11. Significantly enriched proteins of pulldown with mESC and mSP1probe

Table S12. mESC readers_Gene Ontology Terms

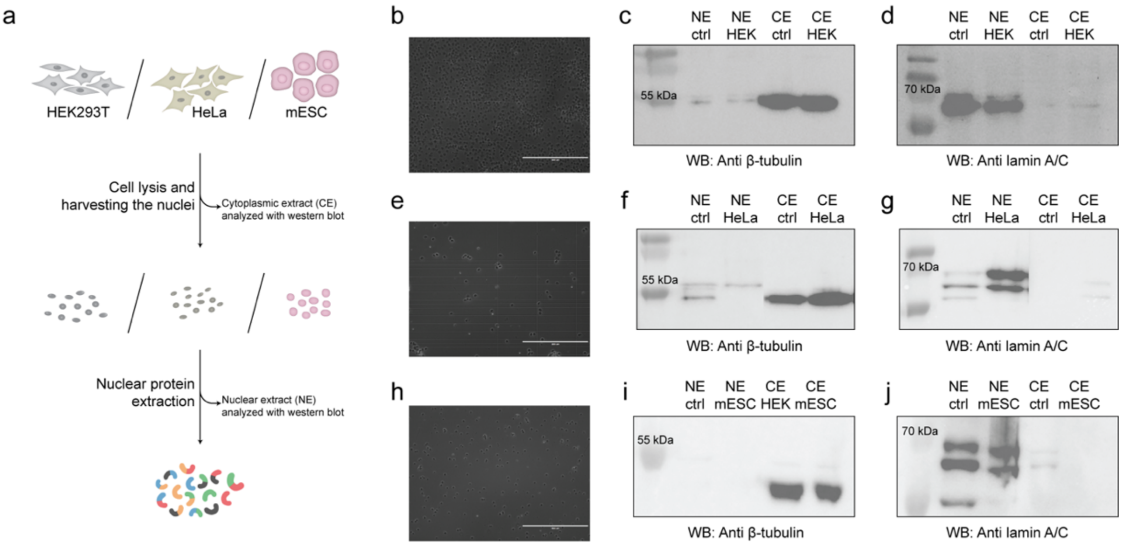


**Supplementary Figure 1. Nuclear protein extraction.** (a) Schematic representation of the nuclear extraction workflow. Cells (HEK293T or HeLa or mESC) were lysed and the nuclei harvested to extract nuclear proteins. (b) The success of the cell lysis is evaluated for HEK293T. The cytoplasmic (CE) and nuclear extracts (NE) are analysed on western blot showing presence of (c) β-Tubulin and (d) lamin A/C in HEK293T. (e) The success of the cell lysis is evaluated for HeLa. The cytoplasmic (CE) and nuclear extracts (NE) are analysed on western blot showing presence of (f) β-Tubulin and (g) lamin A/C in HeLa. (h) The success of the cell lysis is evaluated for mESC. The cytoplasmic (CE) and nuclear extracts (NE) are analysed on western blot showing presence of (i) β-Tubulin and (j) lamin A/C in mESC.

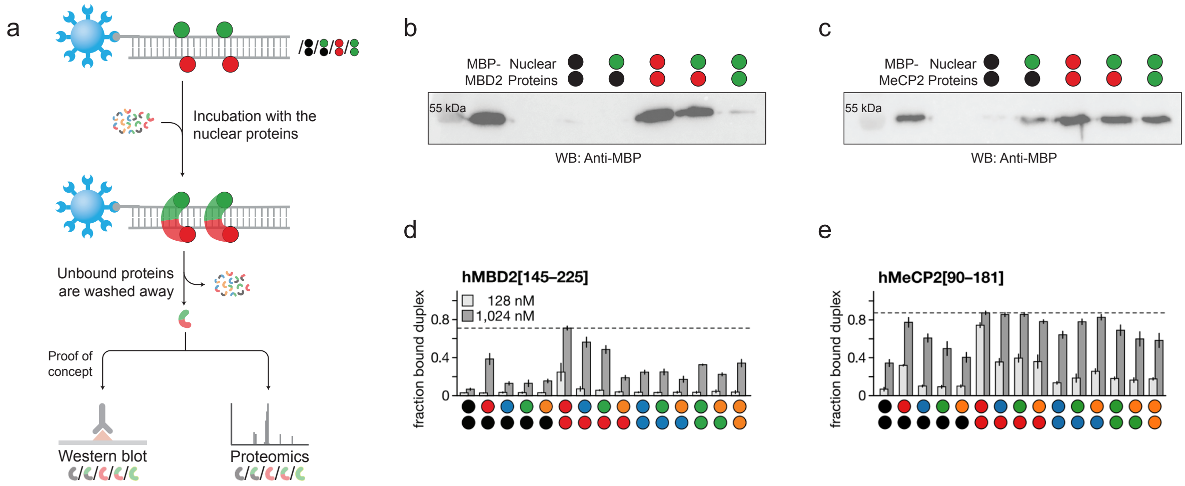


**Supplementary Figure 2. Pulldown based proteomics for discovery of novel readers.** (a) Schematic representation of the pulldown workflow. (b-c) Anti-MBP western blot of the enrichment of (b) MB2-MBP and (c) MeCP2-MBP from nuclear lysates of HEK293, using modified VEGFA probes. (d-e) Reference binding profiles of (d) MBD2 and (e) MeCP2 towards all possible combinations of cytosine modifications based on Buchmuller et al[1]. For colour code, see **Figure 1**.

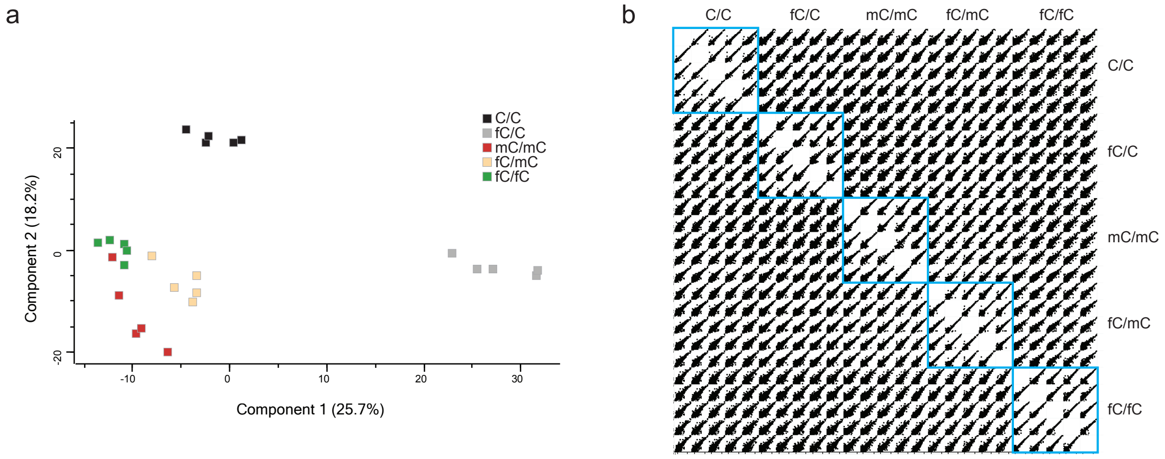


**Supplementary Figure 3. Data correlation and quality control for HEK293T-VEGFA enrichment experiment.** (a) Principal component analysis (PCA) plots of LFQ intensities showing variance across samples. Replicates are shown with the respective colour code per modification as indicated in the legend. (b) Multi-scatter plots of LFQ intensities display pairwise correlations among all technical replicates, grouped by modification condition, with intra-group correlations highlighted in blue squares.

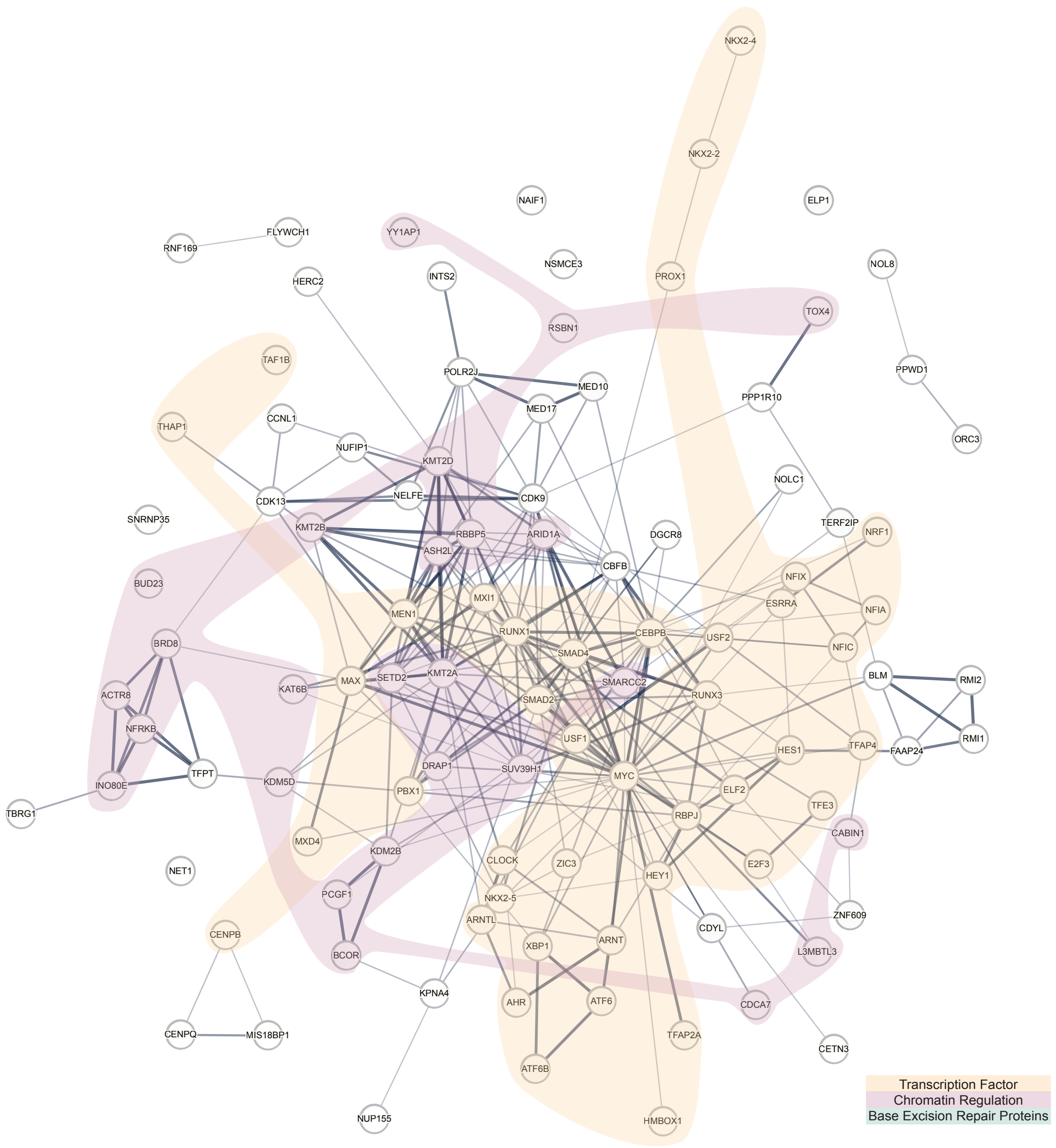


**Supplementary Figure 4**. Interaction network of C/C readers of HEK293T with VEGFA pulldown is shown with annotations for transcription factors in orange, chromatin regulators in purple and base excision repair proteins in green (Analyzed with STRING v12.0[2,3]). For colour code, see **Figure 1**.

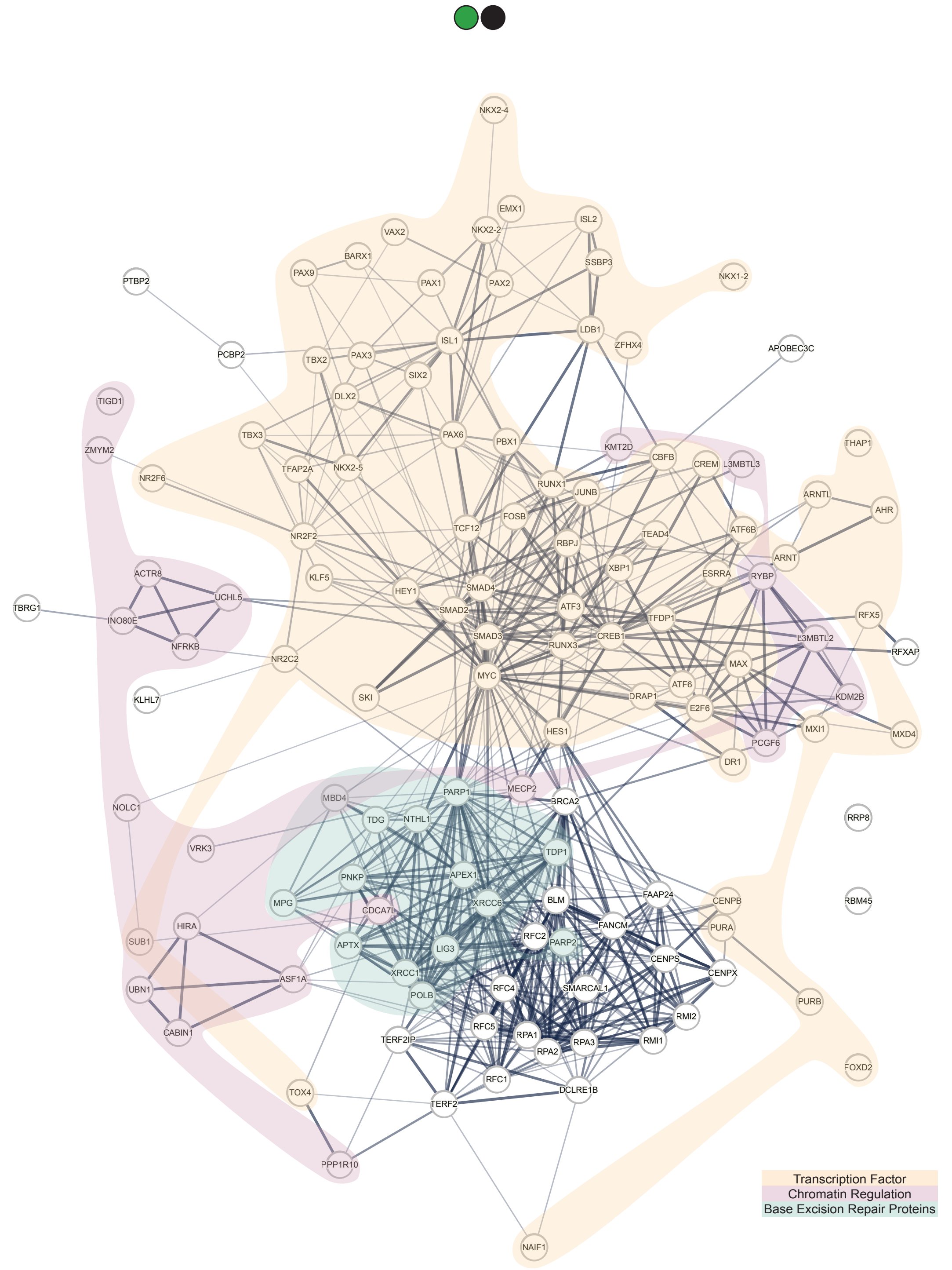


**Supplementary Figure 5**. Interaction network of fC/C readers of HEK293T with VEGFA pulldown is shown with annotations for transcription factors in orange, chromatin regulators in purple and base excision repair proteins in green (Analyzed with STRING v12.0[2,3]). For colour code, see **Figure 1**.

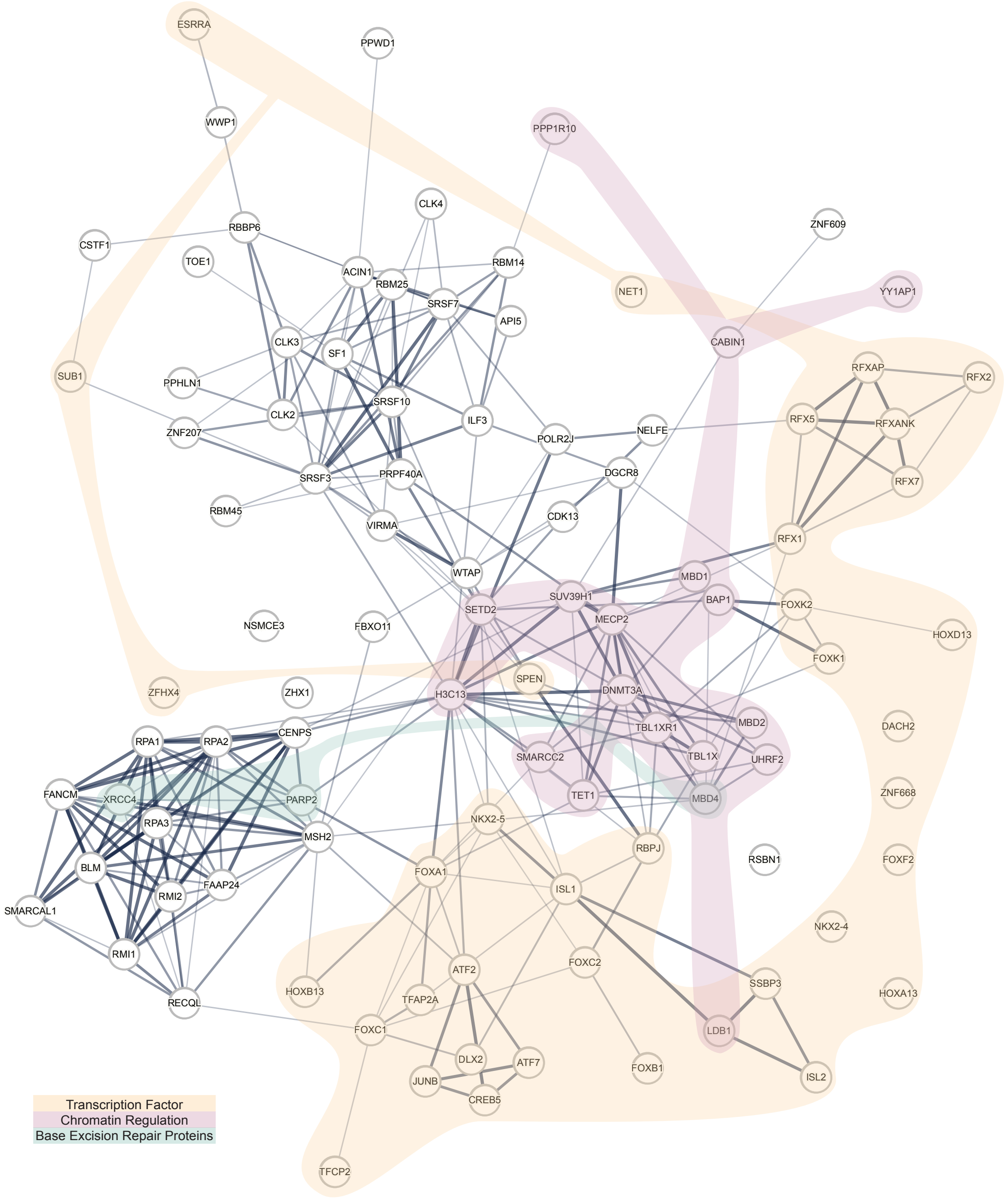


**Supplementary Figure 6**. Interaction network of mC/mC readers of HEK293T with VEGFA pulldown is shown with annotations for transcription factors in orange, chromatin regulators in purple and base excision repair proteins in green (Analyzed with STRING v12.0[2,3]). For colour code, see **Figure 1**.

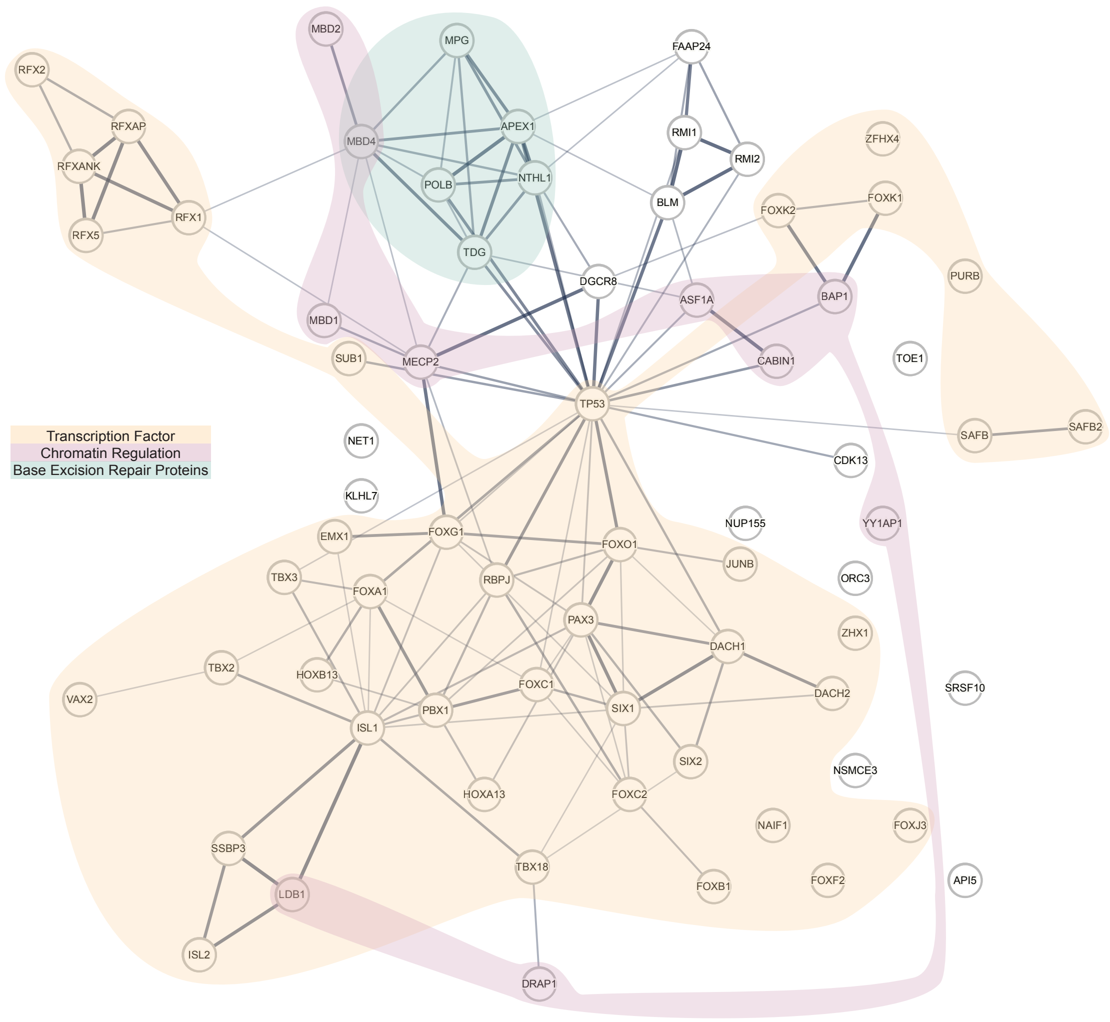


**Supplementary Figure 7**. Interaction network of fC/mC readers of HEK293T with VEGFA pulldown is shown with annotations for transcription factors in orange, chromatin regulators in purple and base excision repair proteins in green (Analyzed with STRING v12.0[2,3]). For colour code, see **Figure 1**.

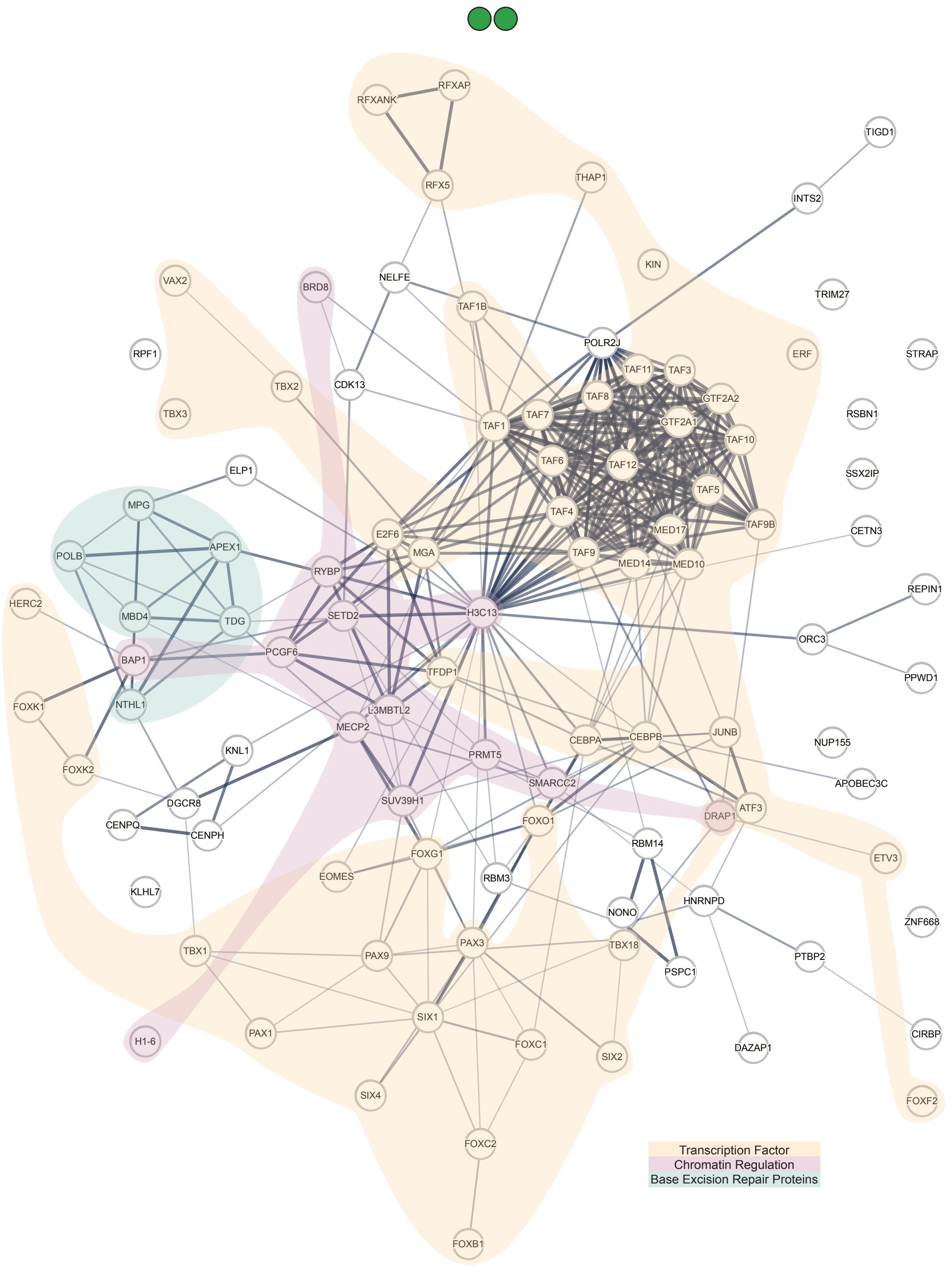


**Supplementary Figure 8**. Interaction network of fC/fC readers of HEK293T with VEGFA pulldown is shown with annotations for transcription factors in orange, chromatin regulators in purple and base excision repair proteins in green (Analyzed with STRING v12.0[2,3]). For colour code, see **Figure 1**.

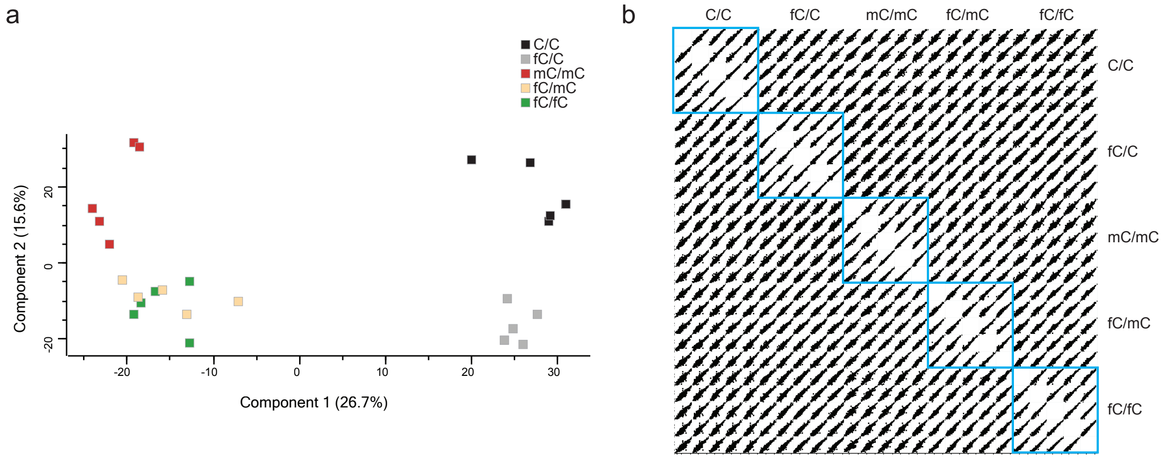


**Supplementary Figure 9. Data correlation and quality control for HEK293T-hSP1 enrichment experiment.** (a) Principal component analysis (PCA) plots of LFQ intensities showing variance across samples. Replicates are shown with the respective colour code per modification as indicated in the legend. (b) Multi-scatter plots of LFQ intensities display pairwise correlations among all technical replicates, grouped by modification condition, with intra-group correlations highlighted in blue squares.

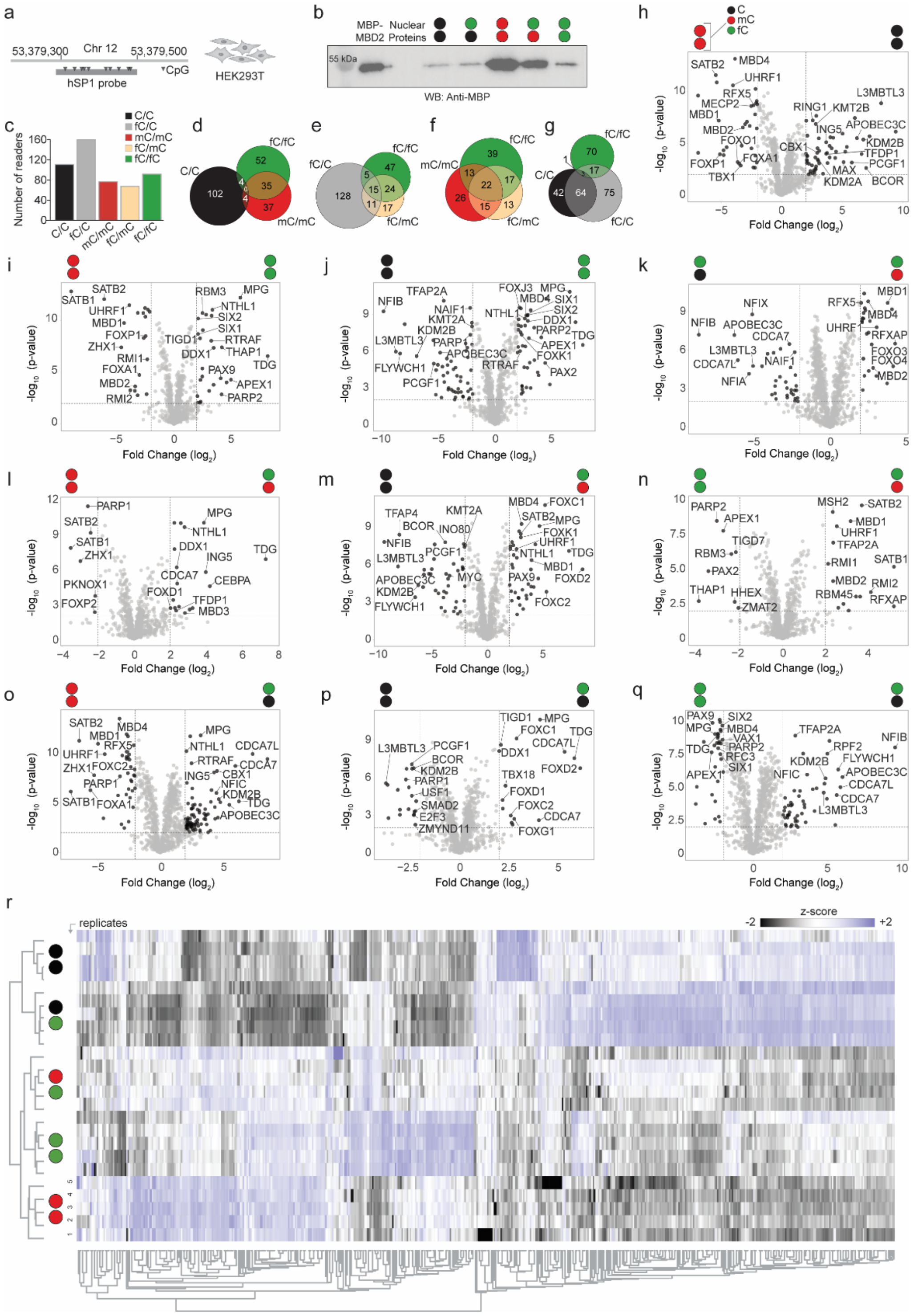


**Supplementary Figure 10**. **Discovery of human readers and antireaders of symmetric and asymmetric fC-modifications by proteomics.** (a) Employed probe design **(hSP1)** and nuclear extract **(HEK293T)** used in this experiment. (b) Anti-MBP western blot of the enrichment of MBD2-MBP spike-in from nuclear lysate using hSP1 probes modified as indicated. (c) Overall number of enriched reader proteins for indicated modifications. (d-g) Venn diagrams showing the overlaps of significantly enriched proteins. Note that Figure 2c-g use a color code describing both DNA strands at once. (h-q) Volcano plots from pairwise comparisons (log2-fold changes > 2 and p-value < 0.01; see Table SI 8 for details). (r) Proteins were filtered to retain only if at least 4 replicates per condition contained a valid LFQ value. ANOVA (permutation-based FDR 0.01) was performed, and significant proteins were z-scored and visualized in a heat map. Missing values were imputed from a normal distribution (downshift 1.8, width 0.3). For colour code, see **Figure 1**.

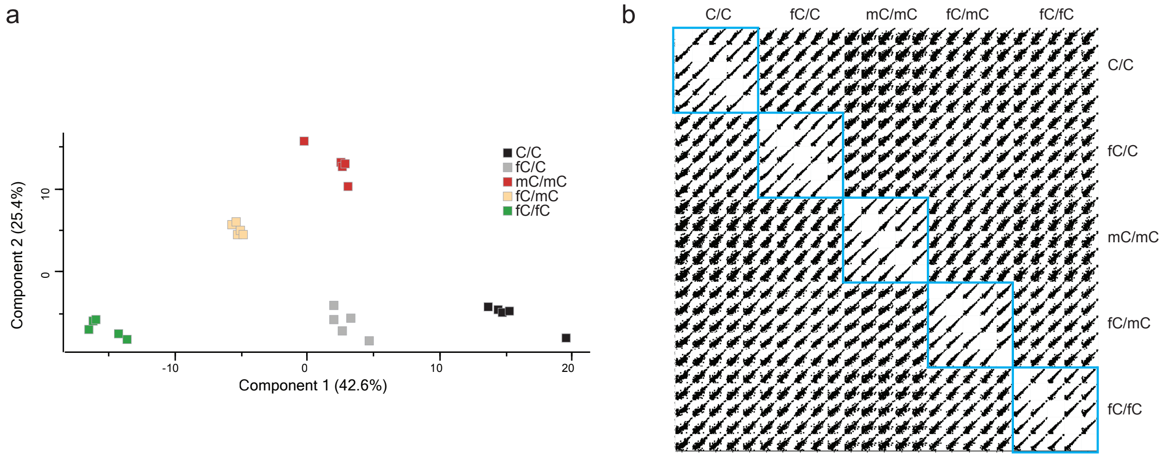


**Supplementary Figure 11. Data correlation and quality control for HeLa-VEGFA enrichment experiment.** (a) Principal component analysis (PCA) plots of LFQ intensities showing variance across samples. Replicates are shown with the respective colour code per modification as indicated in the legend. (b) Multi-scatter plots of LFQ intensities display pairwise correlations among all technical replicates, grouped by modification condition, with intra-group correlations highlighted in blue squares.

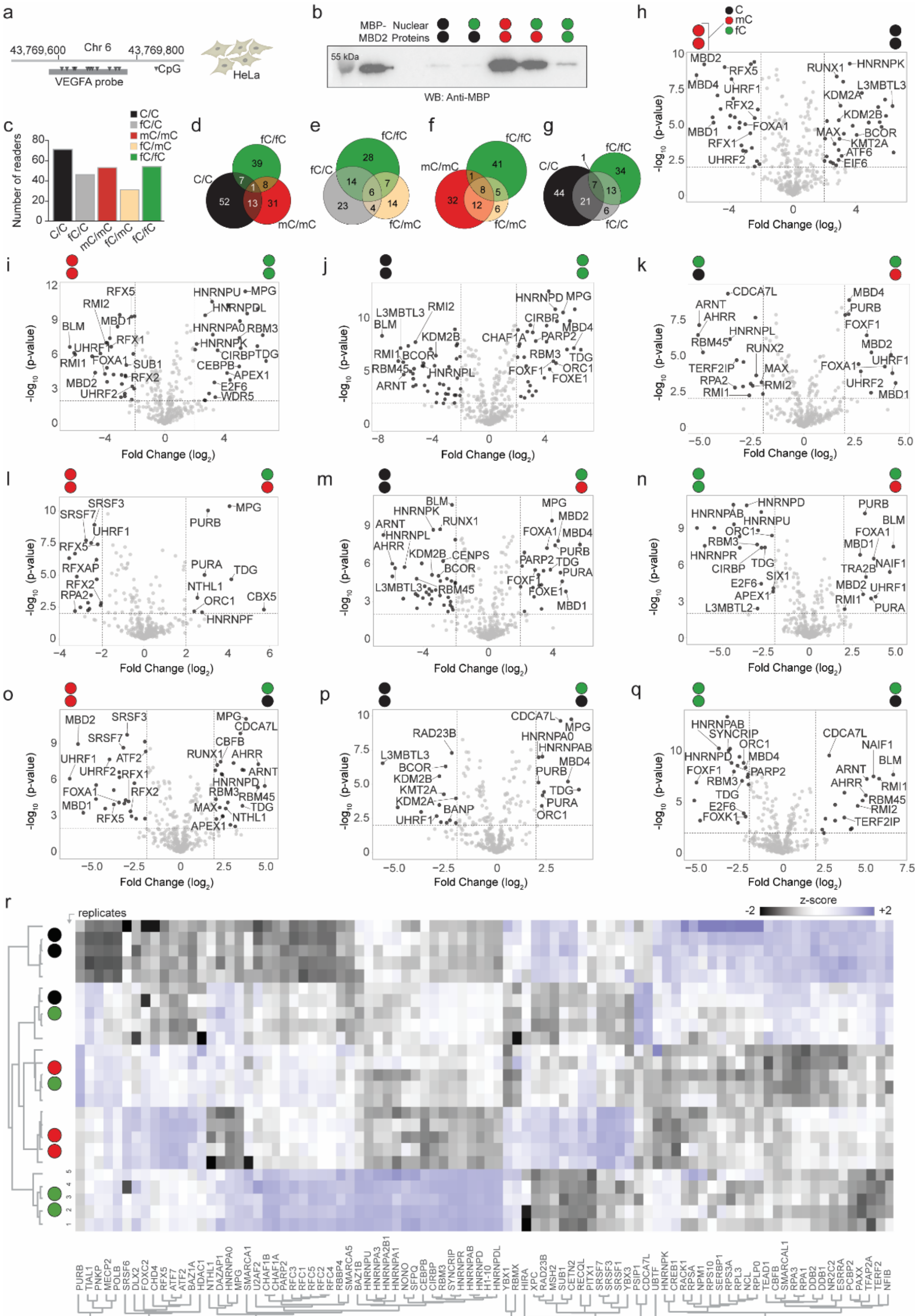


**Supplementary Figure 12**. **Discovery of human readers and antireaders of symmetric and asymmetric fC-modifications by proteomics.** (a) Employed probe design **(VEGFA)** and nuclear extract **(HeLa)** used in this experiment. (b) Anti-MBP western blot of the enrichment of MBD2-MBP spike-in from nuclear lysate using hSP1 probes modified as indicated. (c) Overall number of enriched reader proteins for indicated modifications. (d-g) Venn diagrams showing the overlaps of significantly enriched proteins. Note that Figure 2c-g use a color code describing both DNA strands at once. (h-q) Volcano plots from pairwise comparisons (log2-fold changes > 2 and p-value < 0.01; see Table SI 9 for details). (r) Proteins were filtered to retain only if at least 4 replicates per condition contained a valid LFQ value. ANOVA (permutation-based FDR 0.01) was performed, and significant proteins were z-scored and visualized in a heat map. Missing values were imputed from a normal distribution (downshift 1.8, width 0.3).. For color code, see Figure 1.

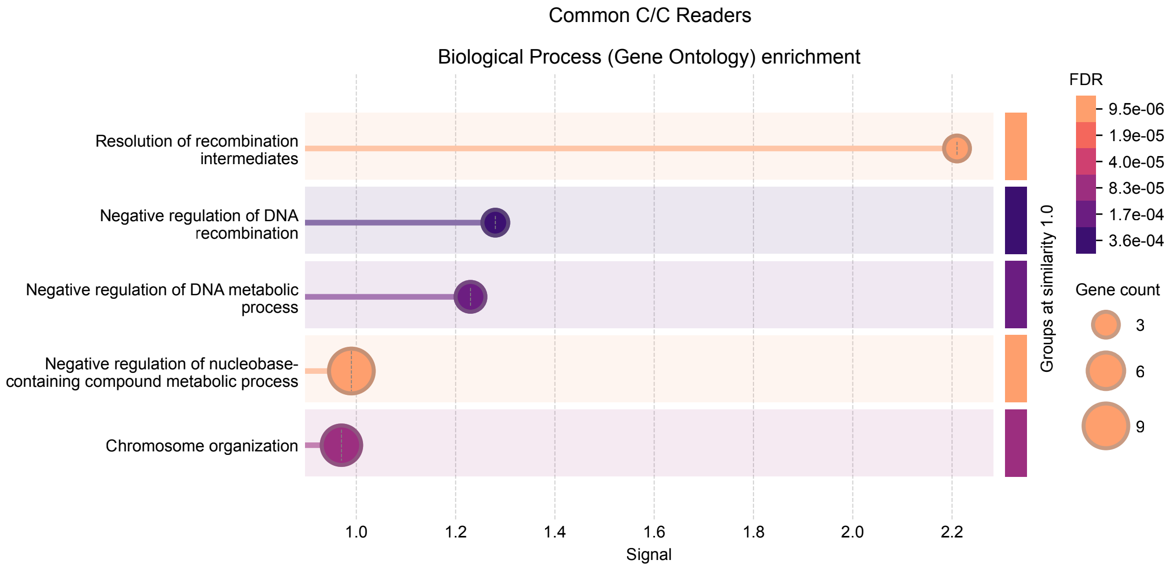


**Supplementary Figure 13**. Gene ontology (GO) [4,5] enrichment analysis of the **common C/C readers,** performed using the STRING database [2,3], revealing significantly enriched biological process terms. Bubble size and color gradient reflect the number of proteins mapped to each term and the statistical significance of enrichment, respectively.

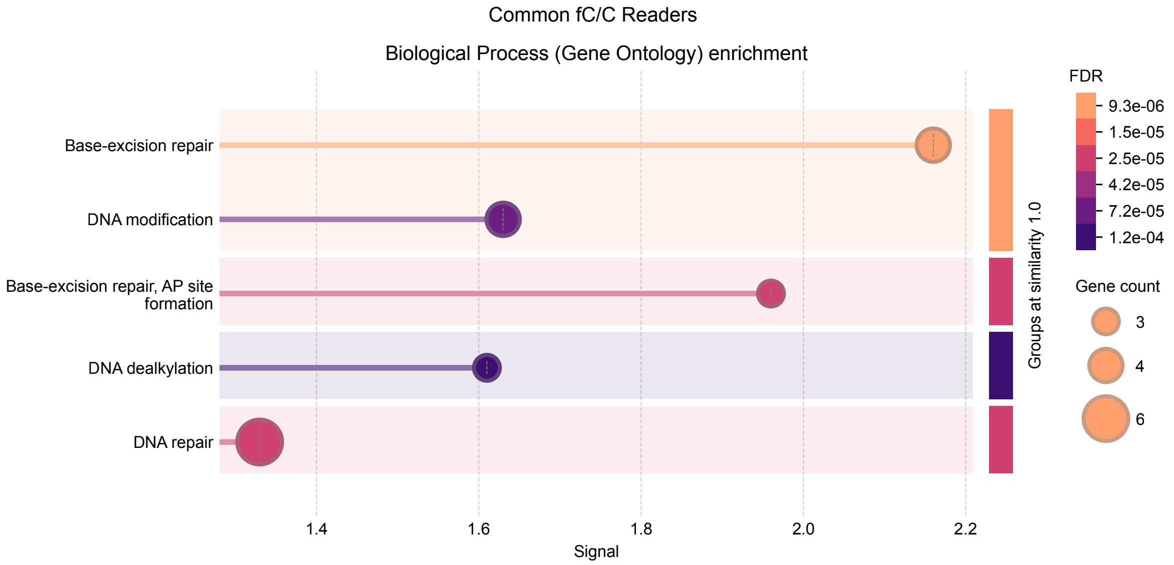


**Supplementary Figure 14**. Gene ontology (GO) [4,5] enrichment analysis of the **common fC/C readers,** performed using the STRING database [2,3], revealing significantly enriched biological process terms. Bubble size and color gradient reflect the number of proteins mapped to each term and the statistical significance of enrichment, respectively.

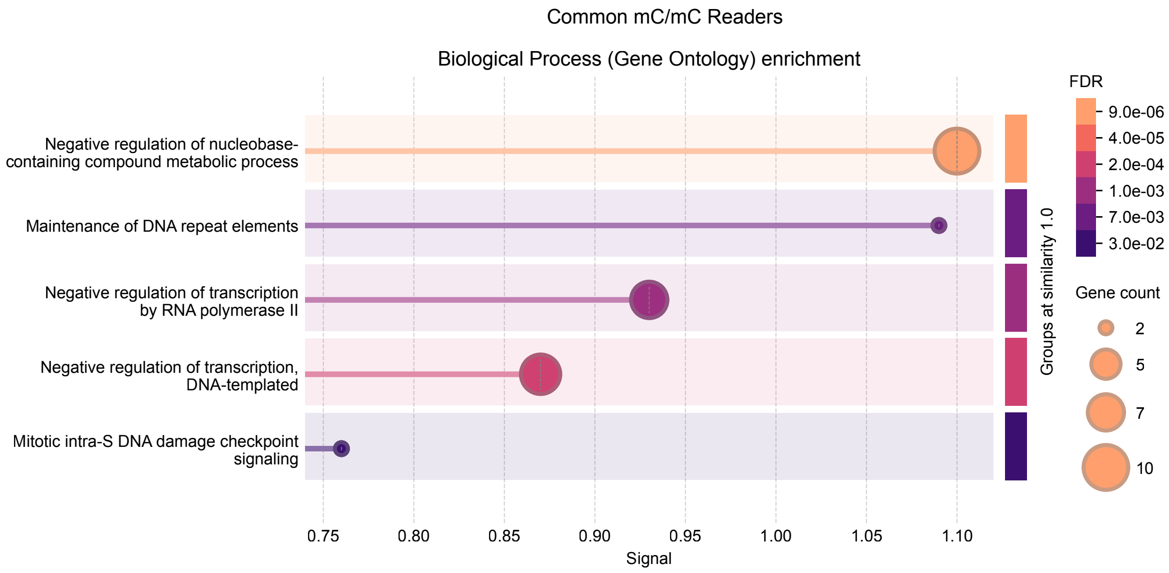


**Supplementary Figure 15**. Gene ontology (GO) [4,5] enrichment analysis of the **common mC/mC readers,** performed using the STRING database [2,3], revealing significantly enriched biological process terms. Bubble size and color gradient reflect the number of proteins mapped to each term and the statistical significance of enrichment, respectively.

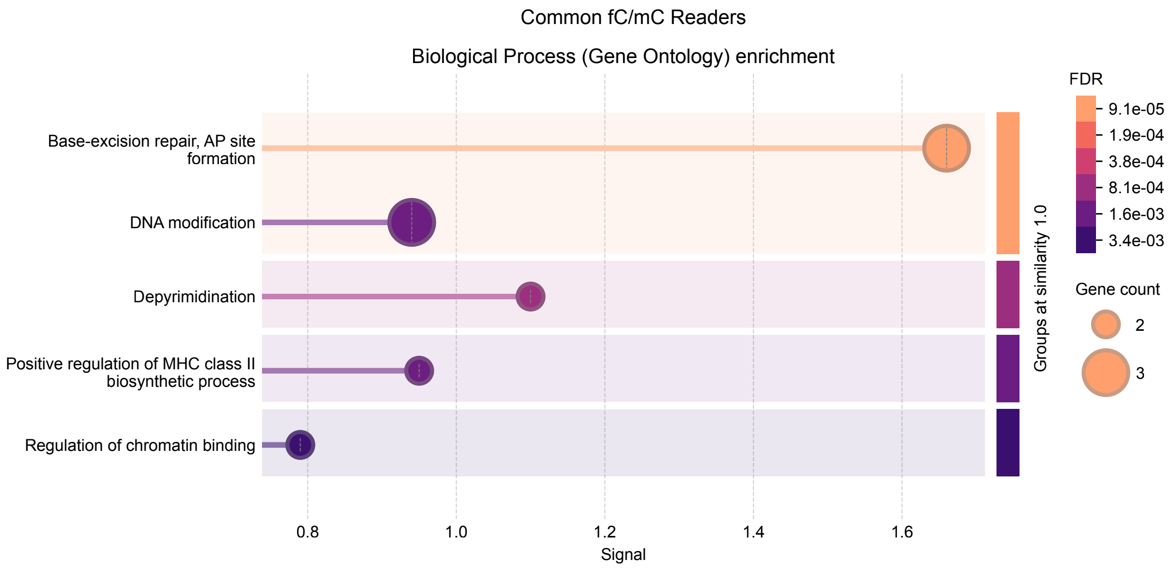


**Supplementary Figure 16**. Gene ontology (GO) [4,5] enrichment analysis of the **common fC/mC readers,** performed using the STRING database [2,3], revealing significantly enriched biological process terms. Bubble size and color gradient reflect the number of proteins mapped to each term and the statistical significance of enrichment, respectively.

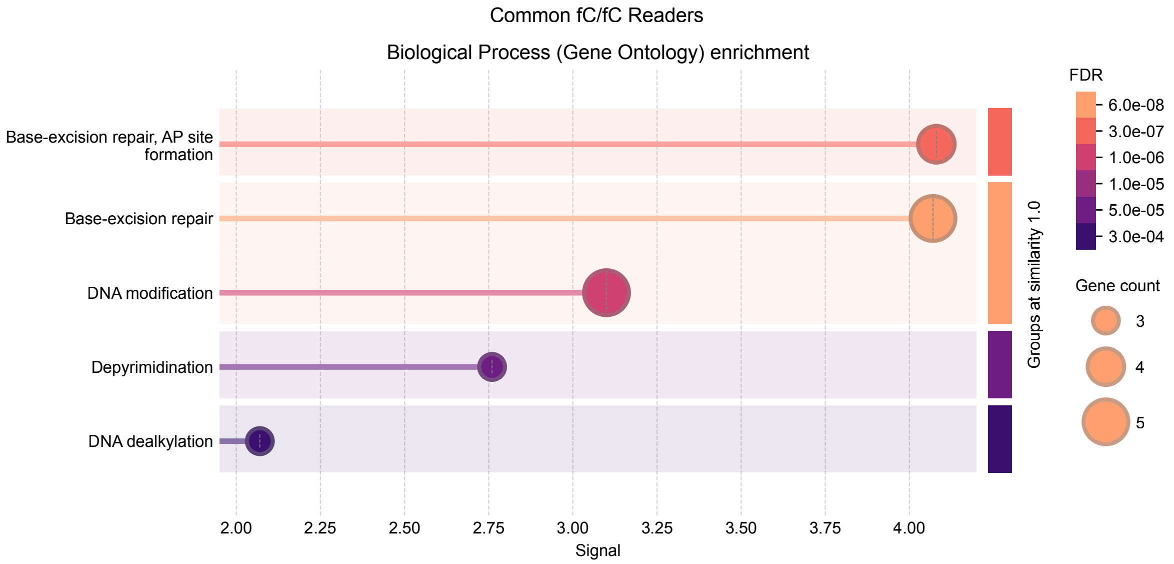


**Supplementary Figure 17**. Gene ontology (GO)[4,5] enrichment analysis of the **common fC/fC readers,** performed using the STRING database[2,3], revealing significantly enriched biological process terms. Bubble size and color gradient reflect the number of proteins mapped to each term and the statistical significance of enrichment, respectively.

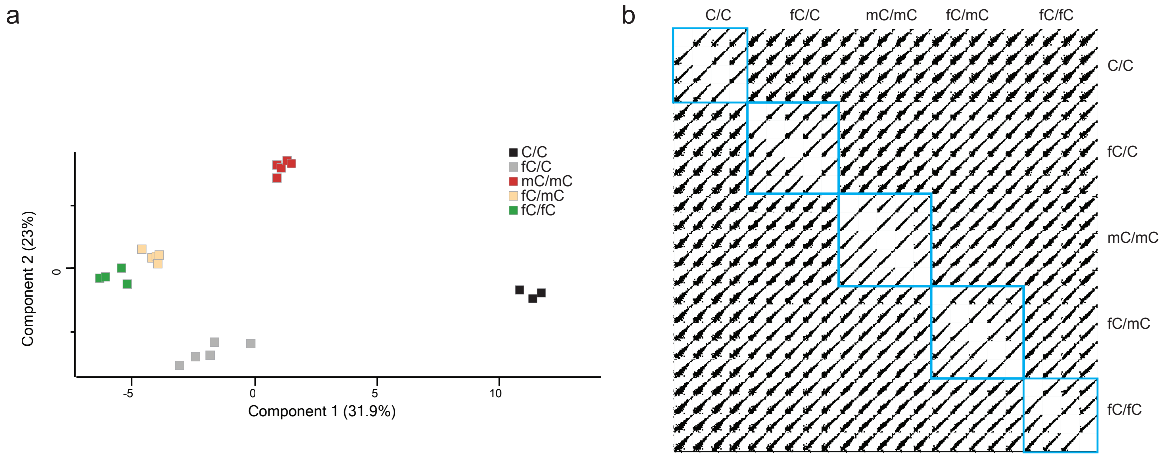


**Supplementary Figure 18. Data correlation and quality control for mESC-mSP1 enrichment experiment**. (a) Principal component analysis (PCA) plots of LFQ intensities showing variance across samples. Replicates are shown with the respective colour code per modification as indicated in the legend. (b) Multi-scatter plots of LFQ intensities display pairwise correlations among all technical replicates, grouped by modification condition, with intra-group correlations highlighted in blue squares.

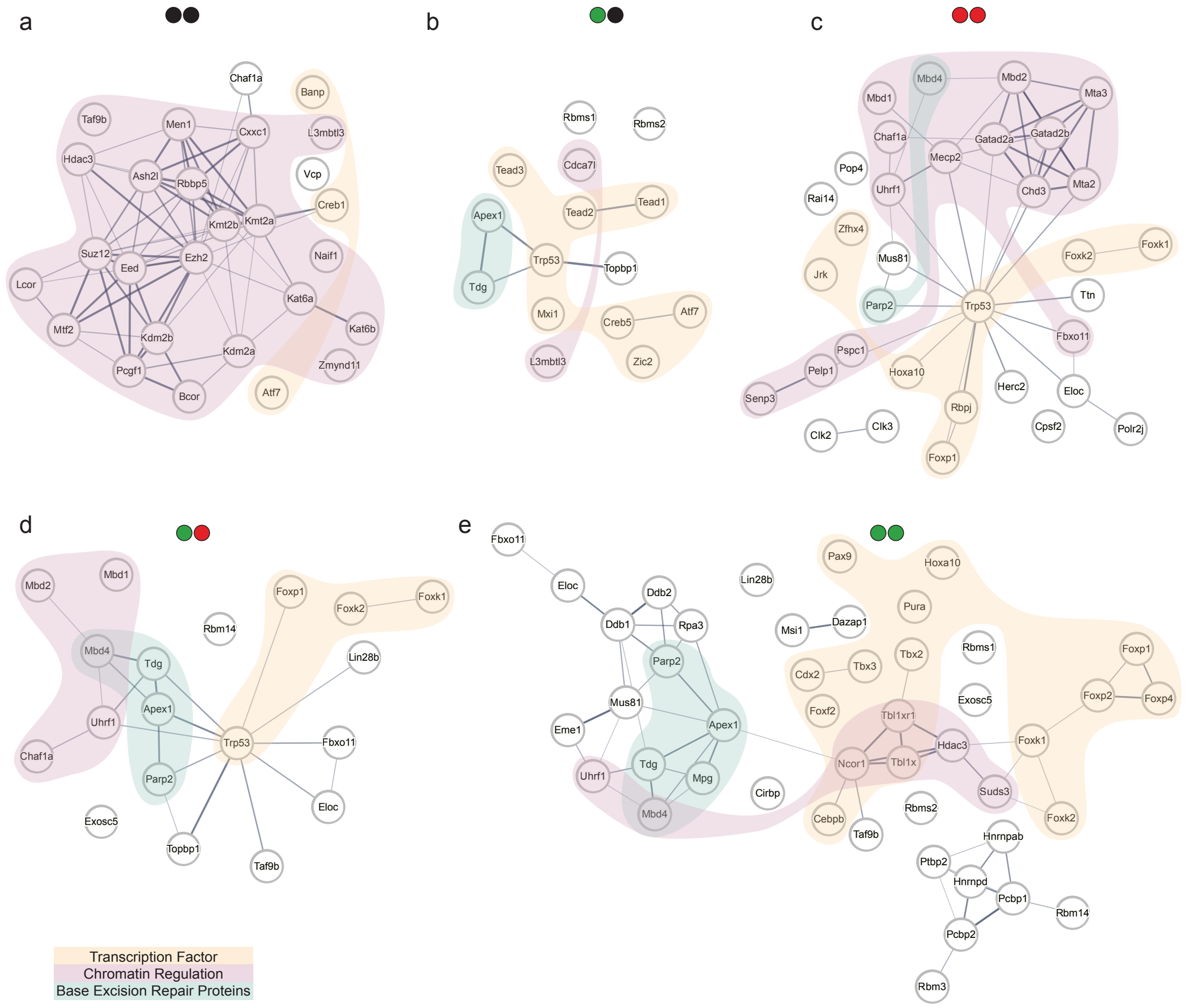


**Supplementary Figure 19.** Interaction network of mESC-mSP1 reader groups with colour code shown on the bottom left. Analysed with STRING database [2,3]. For cytosine modification color code, see Figure 1.

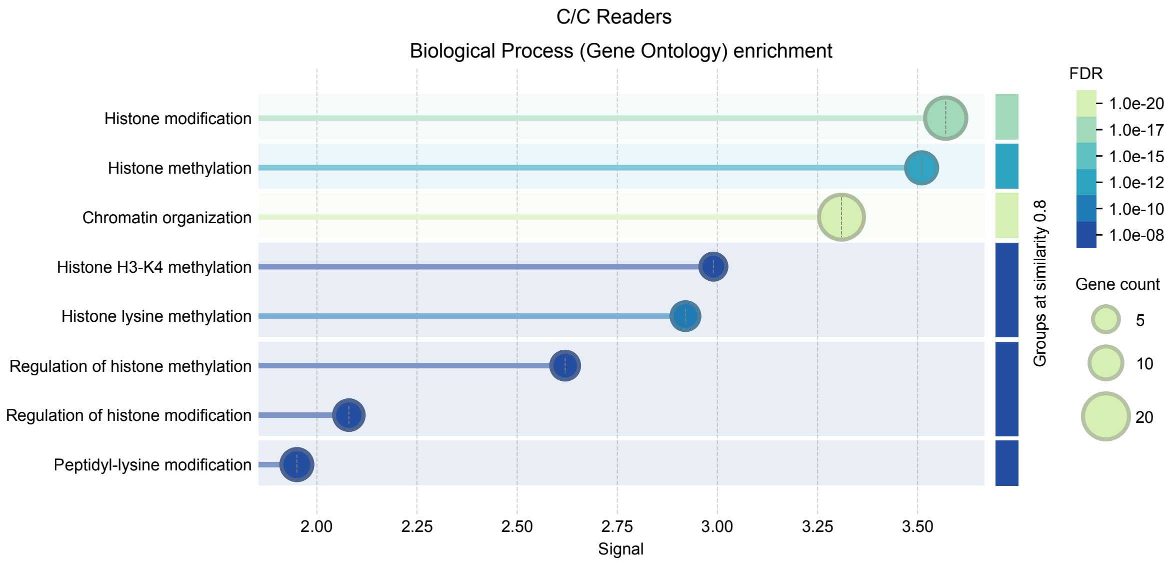


**Supplementary Figure 20**. Gene ontology (GO)[4,5] enrichment analysis of the **mESC C/C readers,** performed using the STRING database[2,3], revealing significantly enriched biological process terms. Bubble size and color gradient reflect the number of proteins mapped to each term and the statistical significance of enrichment, respectively.

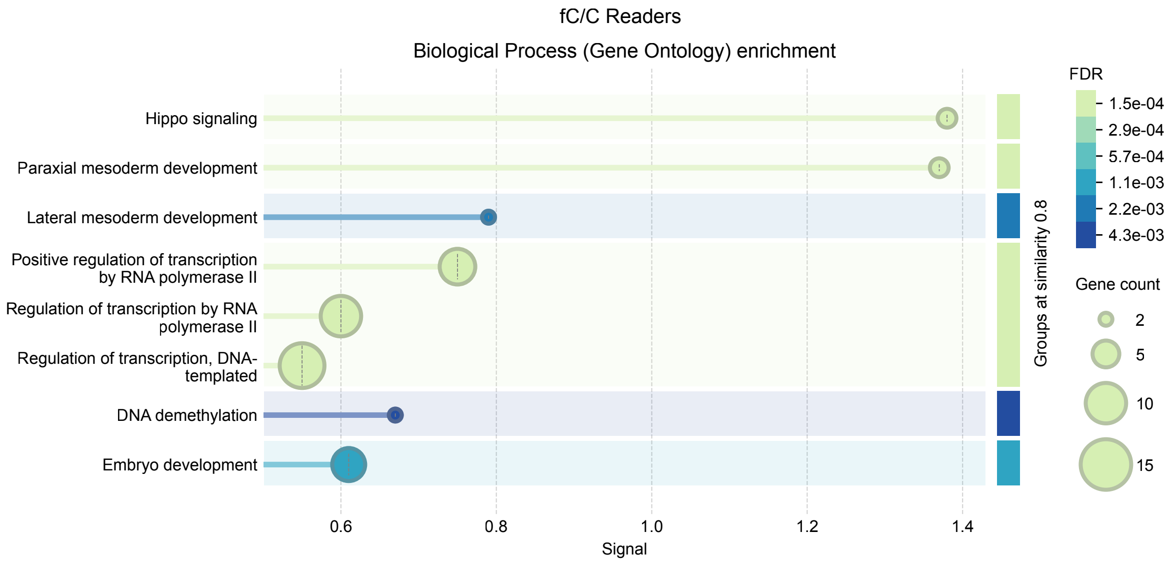


**Supplementary Figure 21**. Gene ontology (GO)[4,5] enrichment analysis of the **mESC fC/C readers,** performed using the STRING database[2,3], revealing significantly enriched biological process terms. Bubble size and color gradient reflect the number of proteins mapped to each term and the statistical significance of enrichment, respectively.

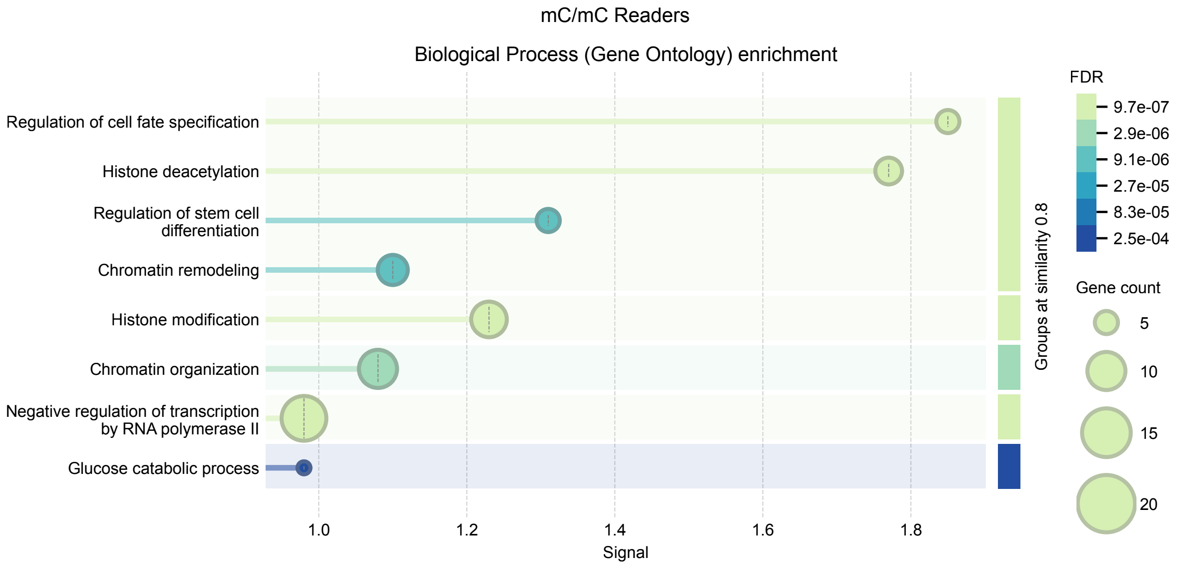


**Supplementary Figure 22**. Gene ontology (GO) [4,5] enrichment analysis of the **mESC mC/mC readers,** performed using the STRING database [2,3], revealing significantly enriched biological process terms. Bubble size and color gradient reflect the number of proteins mapped to each term and the statistical significance of enrichment, respectively.

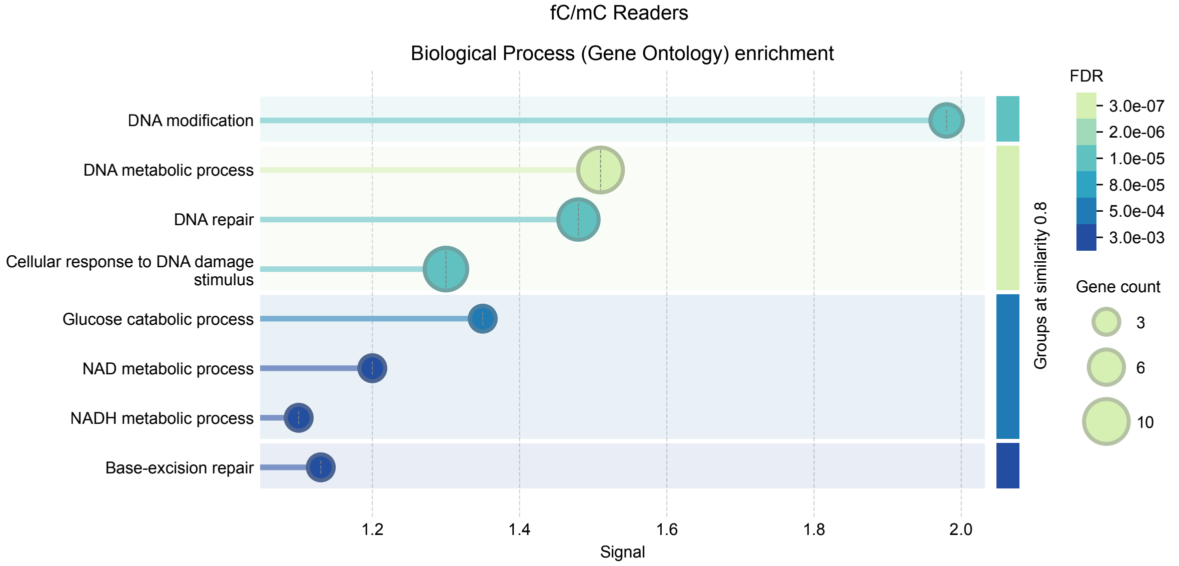


**Supplementary Figure 23**. Gene ontology (GO) [4,5] enrichment analysis of the **mESC fC/mC readers,** performed using the STRING database [2,3], revealing significantly enriched biological process terms. Bubble size and color gradient reflect the number of proteins mapped to each term and the statistical significance of enrichment, respectively.

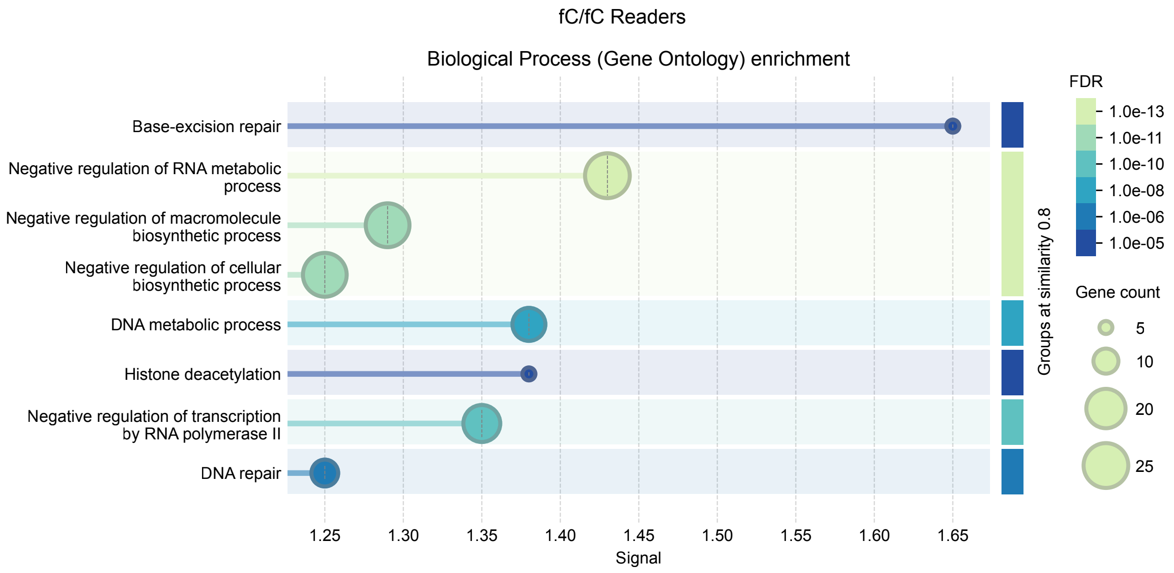


**Supplementary Figure 24**. Gene ontology (GO) [4,5] enrichment analysis of the **mESC fC/fC readers,** performed using the STRING database [2,3], revealing significantly enriched biological process terms. Bubble size and color gradient reflect the number of proteins mapped to each term and the statistical significance of enrichment, respectively.

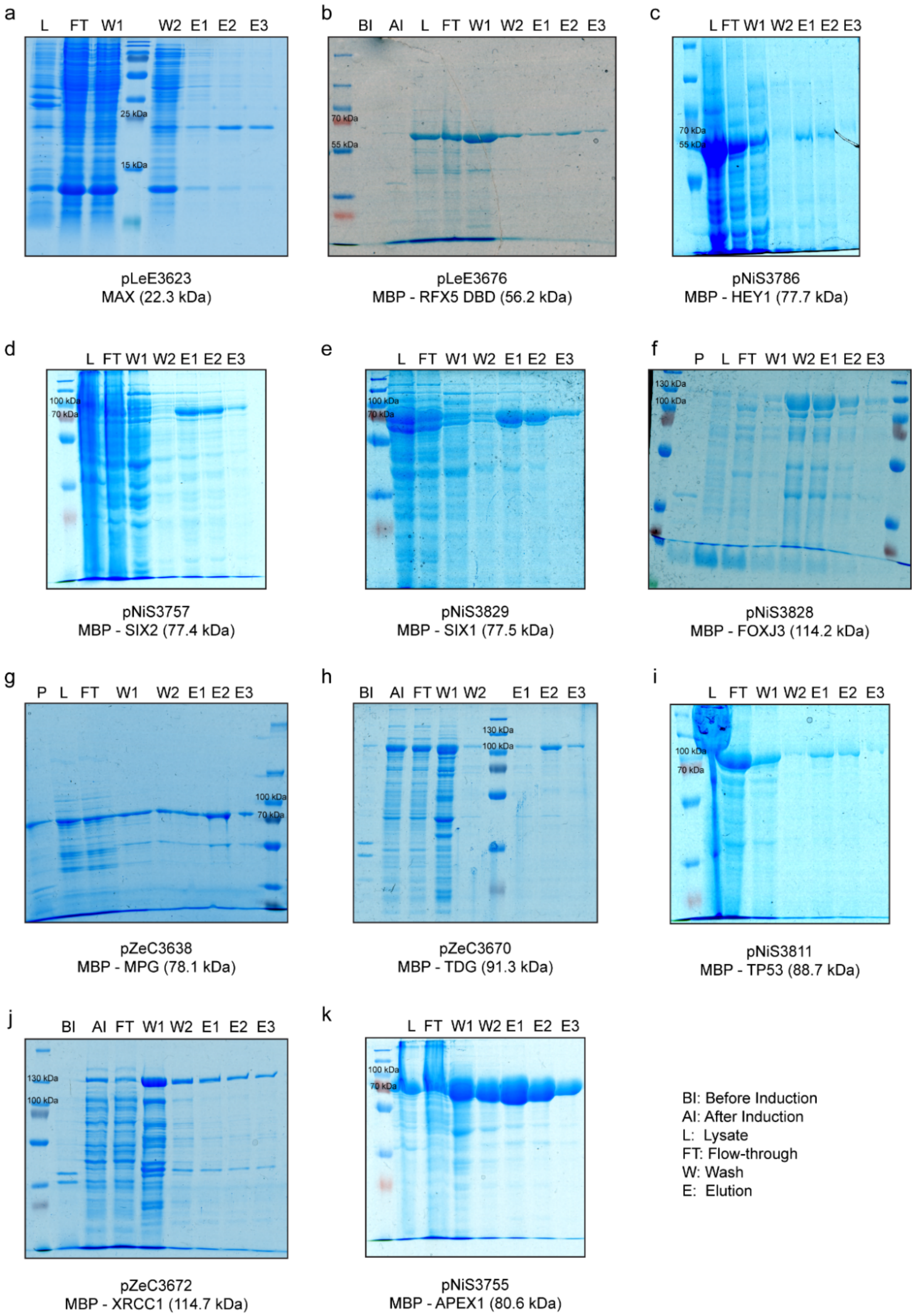


**Supplementary Figure 25**. **Expression and purification of readers for characterization assays.** Coomassie stained 12 % SDS PAGES of purification of (a) MAX, (b) MBP-RFX5 DNA Binding Domain (DBD), (c) MBP-HEY1, (d) MBP-SIX2, (e) MBP-SIX1, (f) MBP-FOXJ3, (g) MBP-MPG, (h) MBP-TDG, (i) MBP-TP53, (j) MBP-XRCC1 and (k) MBP-APEX1 in *E. coli BL21 DE(3) Gold* showing the following fractions: before induction (BI), after induction with 1 mM IPTG (AI), pellet (P) and soluble fractions (L) after sonication referred to as lysate, flow-through (FT) collected after binding to the Ni-NTA resin, wash (W) and elution (E).

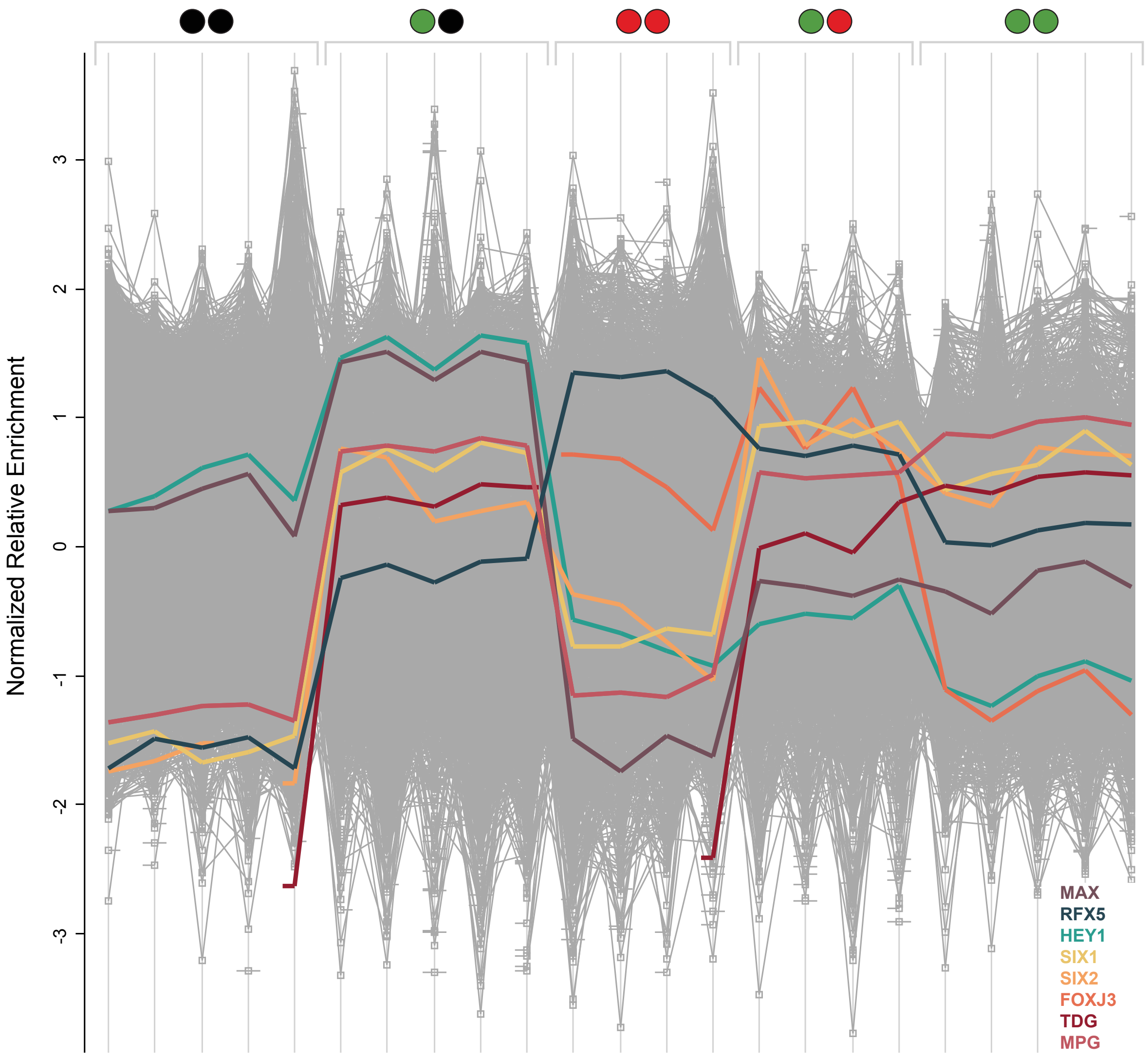


**Supplementary Figure 26**. Binding profiles of the reader proteins from the proteomics experiment employed HEK293T nuclear extracts and VEGFA probe, illustrating the normalized relative enrichment of signal intensities across different CpG modifications and replicates. Each protein profile (MAX, RFX5, HEY1, SIX1, SIX2, FOXJ3, TDG, MPG) corresponds to a distinct relative enrichment and consistency within replicates, enabling direct visual comparison of enrichment patterns.

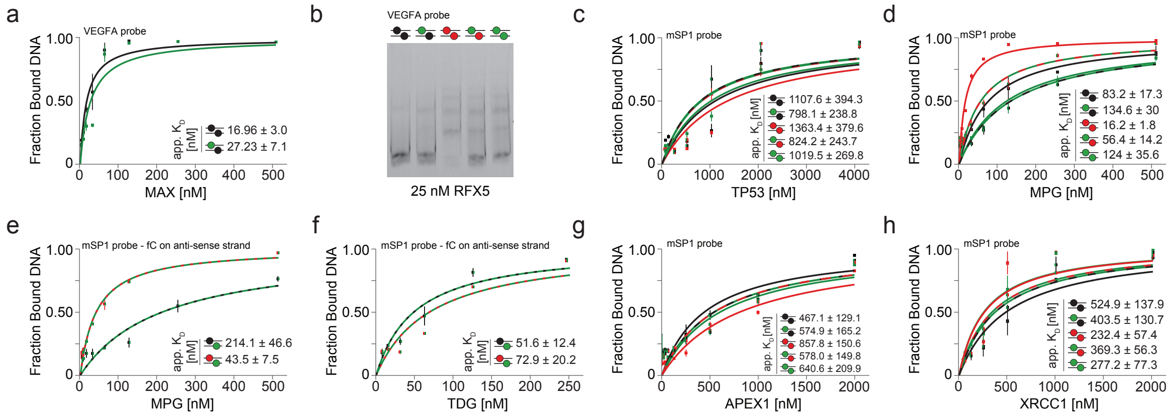


**Supplementary Figure 27**. **Validation of proteins reading fC-modified CpG dyads with distinct dyad symmetry preferences.** (a) EMSA titration experiment with MAX binding to VEGFA probes. (b) EMSA profile of RFX5 binding to VEGFA probes. (c) EMSA titration experiment with TP53 binding to mSP1 probes. (d-e) EMSA titration experiment with MPG binding to mSP1 probes, with fC on sense (d) or antisense (e) strands. (f) EMSA titration experiment with TDG binding to mSP1 probes, with fC on the antisense strand. (g) EMSA titration experiment with APEX1 binding to mSP1 probes. (h) EMSA titration experiment with XRCC1 binding to mSP1 probes. Each experiment was done in duplicates and deviations represented with error bars. For modification colour code, see Figure 1.
